# Genomic functional architecture predicts ecological adaptation across a highly diverse bacterial phylum

**DOI:** 10.1101/2025.07.04.663154

**Authors:** Samuel G. Huete, Killian Coullin, Elodie Chapeaublanc, Rachel Torchet, Nadia Benaroudj, Mathieu Picardeau

## Abstract

Understanding the genetic basis of ecological adaptation is a fundamental challenge of evolutionary biology. *Spirochaetes* are an ancient bacterial phylum of exceptional ecological breadth, encompassing major human pathogens such as the causative agents of syphilis, Lyme disease, and leptospirosis, alongside free-living species from contrasting environments and recently described species that lack the canonical spiral morphology of the phylum. Here, using a curated dataset of genomes representing all cultivable spirochete species, we show that functional genome architecture (defined as the proportional allocation of coding capacity across biological processes) correlates with ecological lifestyle and phenotypic traits in *Spirochaetes* independently of phylogenetic relationships, with host-dependent lineages from distinct evolutionary origins converging on shared functional profiles. Phylum-wide phylogenomic analyses identified *Brachyspira* as the earliest-diverging lineage, revisiting the evolutionary rooting of the phylum, and establishing that the spiral morphology is ancestral and was lost in a single evolutionary transition also associated with coordinated functional changes. Lastly, ancestral genome reconstruction uncovers the common ancestor as a motile, heterotrophic, and spiral-shaped bacterium, with metabolic and structural features not previously described. Together, these results provide an integrated and functional framework for *Spirochaetes*, illustrating how genome architecture can be used to track the ecological diversification of a widespread bacterial phylum.

## INTRODUCTION

*Spirochaetes* (or *Spirochaetota*) form a coherent and ancient phylogenetic phylum (1) of exceptional ecological breadth, whose diversity of lifestyles offers an opportunity for understanding how closely related organisms adapt to radically different environments. The phylum *Spirochaetes* comprises a single class, *Spirochaetia*, and includes four orders: *Brachyspirales*, *Brevinematales*, *Leptospirales*, and *Spirochaetales*. The order *Spirochaetales* is the largest, containing five families and numerous genera, such as *Borrelia*, *Treponema*, or *Spirochaeta* (2). *Spirochaetes* are widely distributed across diverse environments, occupying a broad range of ecological niches from free-living environmental species to host-associated lineages. Despite their common evolutionary origin, they inhabit highly contrasting environments, including highly alkaline lakes (3), the deep subsurface (4), or the gut microbiome (5,6). Members of this phylum display substantial metabolic diversity, existing as anaerobes, facultative anaerobes, microaerophiles, or aerobes. Several genera are of major medical and ecological importance, encompassing both pathogenic and non-pathogenic representatives. Other genera include symbiotic spirochetes inhabiting the hindguts of termites and wood-eating cockroaches (7). Among the human pathogens, *Borrelia* spp. include the agents of Lyme disease (*B. burgdorferi*), one of the most prevalent vector-borne infections in North America and Europe, affecting several hundred thousand people annually (8–10)*. Treponema* spp. include *T. pallidum*, the causative agent of over 50 million cases of syphilis worldwide annually (11). Leptospirosis is a re-emergent zoonotic disease caused by pathogenic *Leptospira* species and accounts for approximately 1 million severe cases and 60,000 deaths every year (12). Lastly, novel taxa, some uncultivated, are being described, but their significance remains to be elucidated (13).

Until recently, all spirochetes were believed to share a specific spiral-shaped morphology and utilize endoflagellar systems for motility (14). However, the discovery of new coccoid, non-spiral *Spirochaetes* species that lack the helical or spiral morphology traditionally considered a defining feature of the phylum challenged long-standing paradigms by suggesting that key spirochete traits such as morphology and motility may be evolutionarily labile (5,15–19). Yet these findings remain fragmented across a limited number of lineages and have not been placed within a phylum-wide evolutionary context. Whether non-spiral spirochetes originated from a single evolutionary transition or multiple independent events across the phylum remains unresolved. Addressing these and similar ecological questions is essential to understand the evolutionary origin of spirochete traits and requires a standardized and comprehensive phylum-wide perspective.

The diversity of lifestyles comprised within a single phylum highlights the potential of *Spirochaetes* as a model for investigating how genomic variation underlies ecological specialization. The particular combination of phylogenetic coherence and ecological diversity, provides a powerful framework to investigate how closely related organisms adapt to distinct lifestyles, bridging the gap between broad tree-of-life comparisons and genus-level studies (20). Importantly, despite their clinical relevance and ecological diversity, key aspects of spirochete biology remain underexplored. *Spirochaetes* are classically difficult to study due to their fastidious growth, recalcitrance to genetic manipulation and complex genomes. *B. burgdorferi* possesses one of the most segmented bacterial genomes, with over 20 plasmids (21). *T. pallidum* has a small genome of around 1.1 Mb and, until very recently, it could not be continuously cultured *in vitro*, dramatically hampering progress in genetic manipulation (22,23). Another characteristic feature is the presence of two circular chromosomes in *Leptospira* spp. (24).

More generally, a central question is to what extent genomic architecture predicts ecological strategies and phenotypic diversification across evolutionary scales. While phylogenetic relationships provide a framework for evolutionary history, they do not fully capture functional and ecological diversification, particularly in lineages that have undergone rapid adaptation or gene content shifts. Therefore, understanding how genome composition maps onto ecological specialization within large bacterial lineages is central to interpreting evolutionary transitions in microbiology, and the *Spirochaetes* phylum provides an exceptional system to address this question directly.

Since the last phylogenetic analysis of the *Spirochaetes* phylum more than a decade ago (2), over 100 new species have been described, yet no study has systematically integrated this expanded diversity in a single reference phylogenetic framework with incorporated functional genomic and trait analyses. In particular, the extent to which functional genome composition reflects ecological strategies across the phylum remains unclear. Here, we address this gap by integrating species-level phylogenomic reconstruction with comprehensive in-depth functional genome analyses across an exhaustive curated dataset of *Spirochaetes* species. We identify lineage-specific functional signatures associated with major ecological lifestyles and show that functional genome composition captures phenotypic and ecological differentiation beyond phylogenetic relationships, revealing convergent ecological strategies across the phylum. Within this framework, we resolve the evolutionary origin of non-spiral spirochetes, demonstrating that the loss of spiral morphology is consistent with a single evolutionary transition associated with coordinated functional changes. Finally, reconstruction of the Last Spirochaetal Common Ancestor (LSCA) features provides insights into the ancestral genomic traits triggering the latter diversification of the phylum. Together, our results establish a unified framework linking genome architecture to phenotypic and ecological transitions across bacterial lineages.

## RESULTS

### Main features of the genomes of Spirochaetes

To construct a comprehensive database of cultured species within the phylum, we selected 172 genomes that ensured both representativeness and highest quality for each species. These genomes are all, except for *Borrreliella chilensis* and *Longinema margulisiae*, RefSeq curated genomes, with an average N50 (the contig length above which 50% of the assembly is contained) of 1.37 Mb, indicating high assembly quality. Among them, 71 are affiliated with *Leptospiraceae*, 39 with *Borreliaceae*, 28 with *Treponemataceae*, 9 with *Brachyspiraceae*, 3 with *Brevinemataceae*, and 22 have unclear taxonomic assignments within *Spirochaetia* (Table S1). In the phylum *Spirochaetes*, genome sizes vary widely, ranging from 0.9 Mb for *Borreliella garinii* (812 proteins) to 4.9 Mb for *Spirochaeta isovalerica* (4318 proteins). Generally, the largest genomes belong to *Leptospiraceae* while the smallest genomes belong to *Borreliaceae*, which is consistent with their vector-borne, host- dependent parasitic lifestyle (Fig 1A, Table S1). Additionally, GC percentage varies significantly, spanning from 24.1% for *Brachyspira intermedia* to 60.9% for *Spirochaeta thermophila*. Genomes from *Brachyspiraceae* and *Borreliaceae* exhibit the lowest GC content, while *Alkalispirochaeta* spp. possess the highest GC contents (Fig 1B, Table S1).

**Figure 1.**
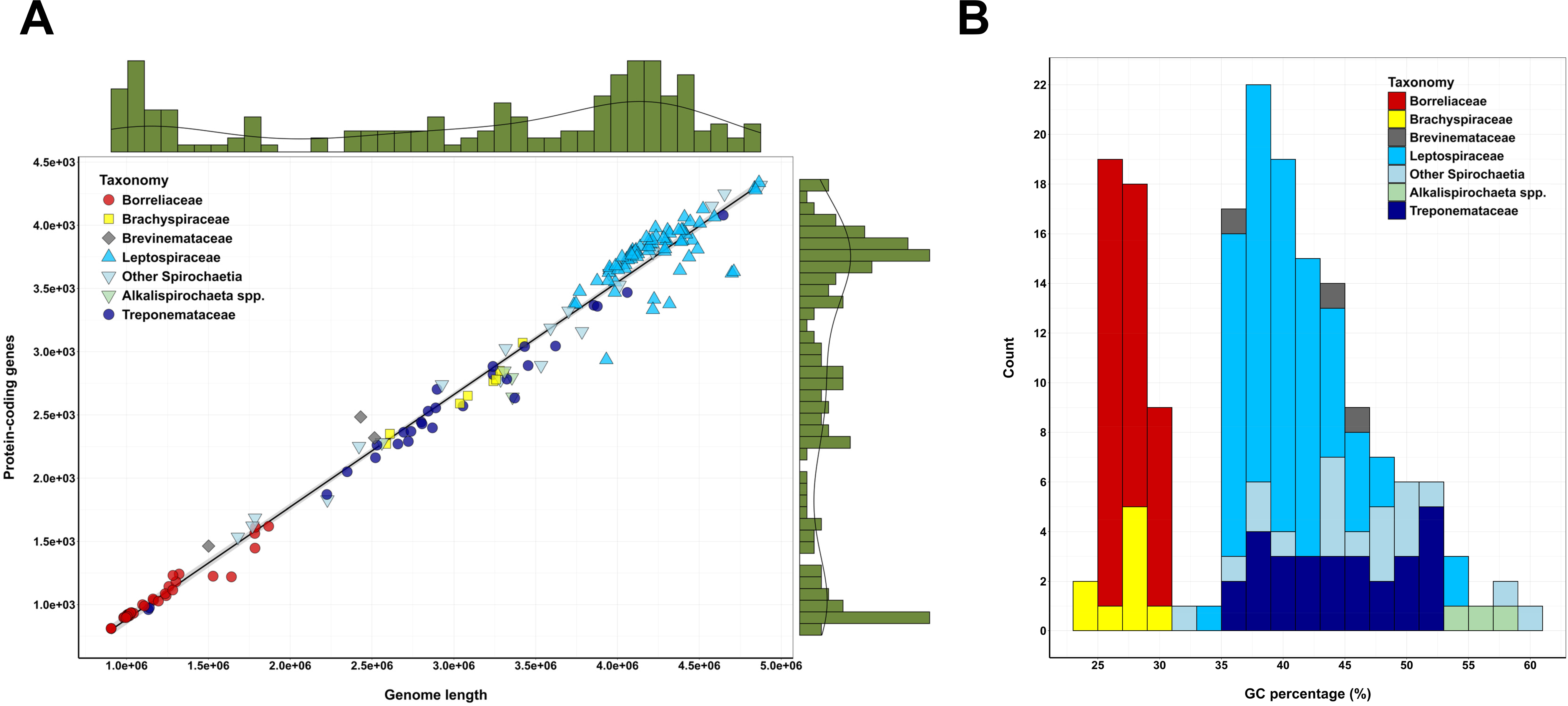
General features of the genomes of *Spirochaetes*. (A) Correlation between genome length (in base pairs) and number of protein-coding genes in the *Spirochaetes* genomes, coloured according to the taxonomy. The green histograms on the sides represent the distribution of the genome length (upper histogram) and number of protein-coding genes (right-side histogram). (B) Distribution of GC content (%) in *Spirochaetes* coloured according to the taxonomy. *Alkalispirochaeta* spp. are separated in both plots for clarity and easy distinction in the text, but taxonomically they must be considered part of *Spirochaetia*.

### A revisited phylogenetic framework for Spirochaetes

To study the evolution of *Spirochaetes*, we inferred orthologs using OMA (Orthologous Matrix), a validated high-accuracy orthology inference framework (25,26), and retrieved 34124 ortholog groups (OGs). Of these, only 1.47% (*N*=503) were shared by at least 50% of the species (*N*=86), highlighting the low conservation of core genes in the phylum (Fig S1). Notably, 50% of the species captured 91% of the reference pangenome (*N*=30906 OGs), illustrating the wide genetic diversity found within this phylum (Fig S1). From these OGs, 140 highly conserved soft-core genome markers were selected for phylogenetic inference. The resulting maximum-likelihood (ML) tree is well resolved under both homogeneous and heterogeneous models of evolution (Fig 2, Fig S2-S3). The topology of the unrooted phylogeny under both models was identical except for the placement of the three *Entomospira* species (Fig S2). In addition, we performed the same process using *Lindowbacteria*, the closest phylum to *Spirochaetes* (27), as a root for both the OG inference and the phylogeny. This time no discrepancies were observed between homogeneous and heterogeneous models (Fig S2). When comparing with the unrooted tree, only the pair *L. stimsonii* and *L. ainazelensis*, two closely phylogenetically related *Leptospira* species of the P1 subclade, showed different placement (Fig S3). Lastly, as an additional control of the topology, we performed a coalescence-based method that handles incomplete lineage sorting using the same soft-core markers. We found more differences between tree topologies when comparing with this method, the major being the placement of the non-spiral *Spirochaetes* species (*Bullifex porci*, *Parasphaerochaeta coccoides*, *Sphaerochaeta pleomorpha*, *Sphaerochaeta globosa* and *Sphaerochaeta halotolerans*) (Fig S4). This difference correlates with the incomplete sorting of some genes in this clade (Fig S4). However, in all phylogenies, the non-spiral *Spirochaetes* form a monophyletic clade, thus providing the first robust phylum-wide evidence that the spiral shape is ancestral to this phylum and was lost only once during the evolution of *Spirochaetes* (Fig S2-S4).

**Figure 2.**
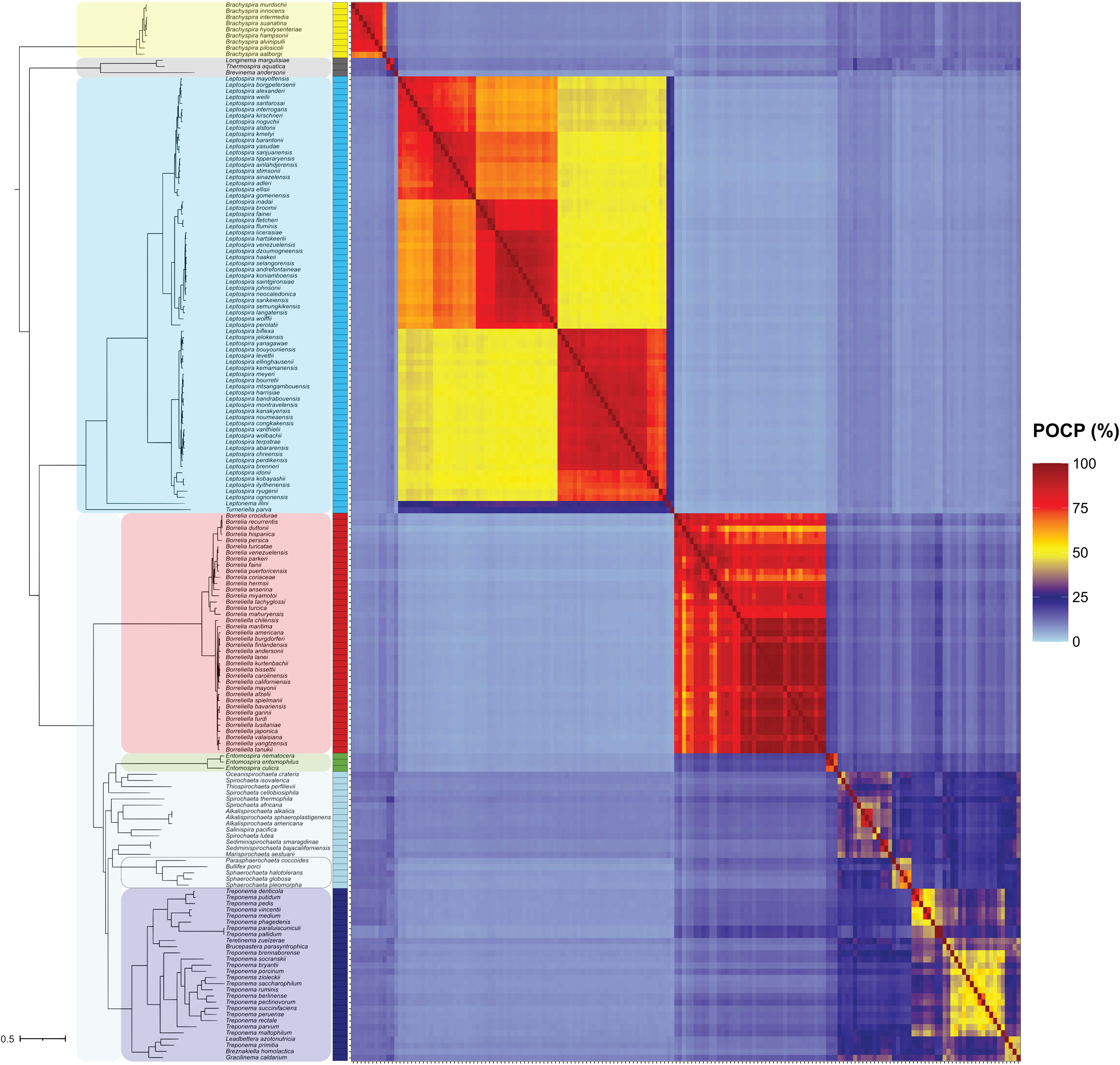
A phylogenetic framework for the *Spirochaetes* phylum. Phylogenetic tree of the cultivable species of *Spirochaetes*, with the main clades highlighted in colours (yellow for *Brachyspira* spp., dark grey for *Brevinematales*, cyan for *Leptospirales*, red for *Borreliaceae*, green for *Entomospira* spp., light blue for intermediate *Spirochaetia*, and dark blue for *Treponemataceae*), represented next to the pairwise matrix of Percentage of Conserved Proteins (POCP, %). The colours in the POCP matrix correspond to the continuous scale represented in the right. The coloured intermediate column represents the main clade classification. The monophyletic clade containing the non-spiral species is also highlighted with a dashed line. This figure is also available on the website <u>spirochase.pasteur.cloud</u> for dynamic exploration.

We thus concluded that the tree topology that best reflects the evolution of the phylum was that obtained under the rooted heterogeneous model (Fig 2). Contrary to previous studies that used more distant phyla to root the phylogeny (2), our results show that *Spirochaetes* separate in two large clades with the first including *Brachyspira* species and the second including the rest of *Spirochaetes* (Fig 2). In this phylogeny, the genus *Brachyspira* is the closest to the root, forming a monophyletic clade. The three sequenced species within *Brevinematales* make a monophyletic clade where *Brevinema andersonii* is the longest branch in that clade. Although *Brachyspira* and *Brevinematales* lineages are only represented by a limited number of genomes, their placement was consistent across all reconstructions and evolutionary models (Fig S2-S4) suggesting a robust evolutionary signal, although further sampling may help refine their exact phylogenetic placement. Similarly, the three only sequenced species within *Entomospira* make a monophyletic clade with a long separation from other *Spirochaetia* species (Fig 2).

As previously described, among the three most represented genera in *Spirochaetes* (*Leptospira* spp., *Borreliaceae* spp., and *Treponema* spp.), *Leptospira* and *Borreliaceae* form monophyletic clades subdivided in four and two major subclades, respectively (Fig 2) (28,29). However, the classification of the clade containing *Treponema* species is less clear. While all *Treponemataceae* species form a monophyletic clade alongside the recently described *Breznakiellaceae* family (30,31), the naming of some of these genera may lead to confusion. Based on this and prior phylogenomic evidence (30,31), we propose reclassifying the species *Treponema primitia* to a different genus and renaming *Teretinema zuelzerae* and *Brucepastera parasyntrophica* to *Treponema zuelzerae* and *Treponema parasyntrophica*, respectively. Such change would imply that termite gut treponemes in the *Breznakiellaceae* family would conform a basal clade to all *Treponema* spp. which would then separate in two main subclades (Fig 2). To further clarify the status of *T. zuelzerae* and *B. parasyntrophica*, we performed another phylogenetic inference restricted to the *Treponematales* clade. This increased by over 3-fold the number of phylogenetic markers (*N*=440), thus representing the most robust phylogenetic inference performed to date in *Treponema* species. Our data confirmed that *Treponema* species subdivide in two main subclades, the first (T1) containing *T. pallidum* and *T. denticola* alongside *T. zuelzerae* and *B. parasyntrophica*, among other treponemes, and the second (T2) containing *T. maltophilum* and *T. porcinum*, among other treponemes (Fig S5) (30,31).

Overall, our phylogenomic analyses support an evolutionary scenario where the Last Spirochaetal Common Ancestor (LSCA) gave rise to two main clades, separating *Brachyspira* species from the rest of *Spirochaetes* early in the phylum’s evolution.

### Pairwise comparisons support the phylogenetic analysis

Pairwise comparison analyses using average nucleotide identity (ANIb, Fig S6), kmer-based tetranucleotide frequencies (TETRA, Fig S7), and the percentage of conserved proteins (POCP, Fig 2), all showed strong concordance with the phylogeny presented in Fig 2. As previously described for other prokaryotes (32), POCP was the most consistent predictor of genus-level boundaries within *Spirochaetes* (Fig S8). All *Brachyspira* spp. and members of *Borreliaceae* share POCP values ≥ 50%, supporting their classification within single, standard cohesive genera. In contrast, some *Leptospira* spp. and *Treponema* spp. share lower intra-genus POCP values below 50%, reflecting a higher degree of genomic and evolutionary divergence than *Brachyspira* and *Borreliaceae* (Fig 2, Fig S8, Table S2).

Within *Treponema*, POCP-based clustering reveals two major groups: the first including a subset of the T1 subclade (*T. denticola*, *T. putidum*, *T. pedis*, *T. vincentii*, and *T. medium*, Fig 2), and the second including all T2-subclade treponemes (Fig 2, Fig S8). Notably, the *Treponema* genus falls below classical genus-level classification thresholds, indicating that, as currently defined, it aggregates lineages with substantially distinct genomic divergence. This inconsistency likely reflects the historical expansion of the genus based on ecological and host-associated traits (33,34) rather than uniform genomic criteria (30,31). We determined that applying a uniform POCP threshold of 45% for genus definition, as supported by the global distribution of values across the phylum (Fig S8), would therefore imply the subdivision of *Treponema* into multiple genera, while conversely supporting the unification of currently separated taxa such as members of the *Borreliaceae* family (Fig S8).

Similar inconsistencies based on POCP clustering are observed in other lineages. Our results suggest that *Longinema margulisiae* and *Thermospira aquatica*, two *Brevinematales*, should be renamed to be included in the same genus. This applies as well to *T. primitia* and *Leadbettera azotonutricia* (Fig 2, Table S2). Together, our phylum-wide results highlight that genus nomenclature within *Spirochaetes* should be treated with caution since it does not accurately reflect uniform levels of genomic divergence for the same taxonomic status.

Altogether, our results support a revised phylogenetic framework of the phylum *Spirochaetes* and show that standardized phylum-wide genomic similarity metrics, such as POCP-based clustering, may help delineate evolutionary relationships and lineage diversification more consistently than current taxonomic nomenclatures. These discordances also indicate that current evolutionary frameworks alone may not fully capture the functional and ecological diversification of the phylum.

### Cluster-based annotation improves functional characterization in Spirochaetes

We next asked whether genome-wide functional information could capture ecological diversification that phylogenetic and genomic similarity metrics alone do not resolve. To characterize the functional repertoire encoded across *Spirochaetes*, we annotated ortholog groups (OGs) using a cluster-based approach with fine-grained orthology (see Methods). This strategy leverages high-confidence OG inference to propagate functional information across related sequences, thereby improving annotation in divergent proteins. Functional coherence within OGs was confirmed by individually annotating each sequence of the 491 most prevalent OGs (63423 proteins, 14.5% of the total dataset), of which 96.3% of the annotated sequences showed consistent COG categories within their OG, thus validating our approach. This method allowed us to assign a COG category to 58% (*N*=19893) of all OGs, with an average 70% annotation level per species (ranging 48–87%) (Fig S9). In addition, we successfully assigned a maximum annotation level to 83% (*N*=28389) of all OGs, with an average 88% annotation level per species (ranging 64–100%) (Fig S9).

Interestingly, *Leptospira* spp. had a significantly higher number of OGs of putative eukaryotic origin than other *Spirochaetes*, including proteins related to actin cytoskeleton dynamics, growth regulation, and transcriptional control (Fig S10, Table S3). Most of these OGs were classified among the functional categories of posttranslational modification (O, 14.2%), signal transduction mechanisms (T, 12.2%), lipid metabolism (I, 8.8%) or unknown function (S, 19.6%) (Table S3). About a third of the eukaryotic OGs (*N*=44, 29.7%) were assigned to the *Metazoa* kingdom, 20.2% (*N*=30) were assigned to the *Fungi* kingdom, and 17.6% (*N*=26) were assigned to the *Streptophyta* phylum (Table S3). Other represented taxonomic levels were the *Aconoidasida* class (3.4%, *N*=5) or the *Opisthokonta* clade (1.4%, *N*=2). Remarkably, some of the assigned functions of these OGs are typically associated with eukaryotic cellular processes and are uncommon in bacterial genomes, suggesting the presence of eukaryotic-like domains or proteins that may have been acquired through horizontal gene transfer or evolved convergently (Table S3) (35,36). Given the ability of *Leptospira* species to colonize diverse hosts and environmental reservoirs, the presence of such eukaryotic-like functions may reflect molecular strategies for interacting with unicellular eukaryotes, niche adaptation or modulating host processes (37).

In contrast, *Borreliaceae* spp. showed a significant enrichment in OGs of viral origin (Fig S10). These elements are consistent with the known abundance of phage-derived plasmids in *Borreliaceae* (21) and suggest that viral sequence integration has been a recurrent feature of genome evolution in this lineage.

### Functional genome composition differentiates ecological and phenotypic structure in Spirochaetes

We next analysed the proportion of each genome assigned to functional categories (COG) and observed distinct functional signatures across the phylum and among different clades (Fig 3A). Principal component analysis (PCA) evidenced that 81% of the total functional variability is captured by two linear components (PC1 and PC2), indicating that a limited number of functional axes structure genome diversity in *Spirochaetes* (Fig S11). These axes are primarily driven by variation in transcription (K), signal transduction (T), energy production (C), and, additionally, secondary metabolism (Q), which contributes the most to PC2 (16.28%) (Fig S11). In contrast, categories such as RNA processing (A) and chromatin structure (B) contributed minimally (1.90-1.96%), due to their low representation across the phylum (Fig S11, Table S4). However, some lineage-specific enrichments were observed, like chromatin-associated functions (B) in *Leptospira* (13-18%) (Fig 3A, Fig S11, Table S4). Together, these results indicate that variation in regulatory and metabolic capacities, rather than housekeeping functions, define the major axes of functional diversification in *Spirochaetes*.

**Figure 3.**
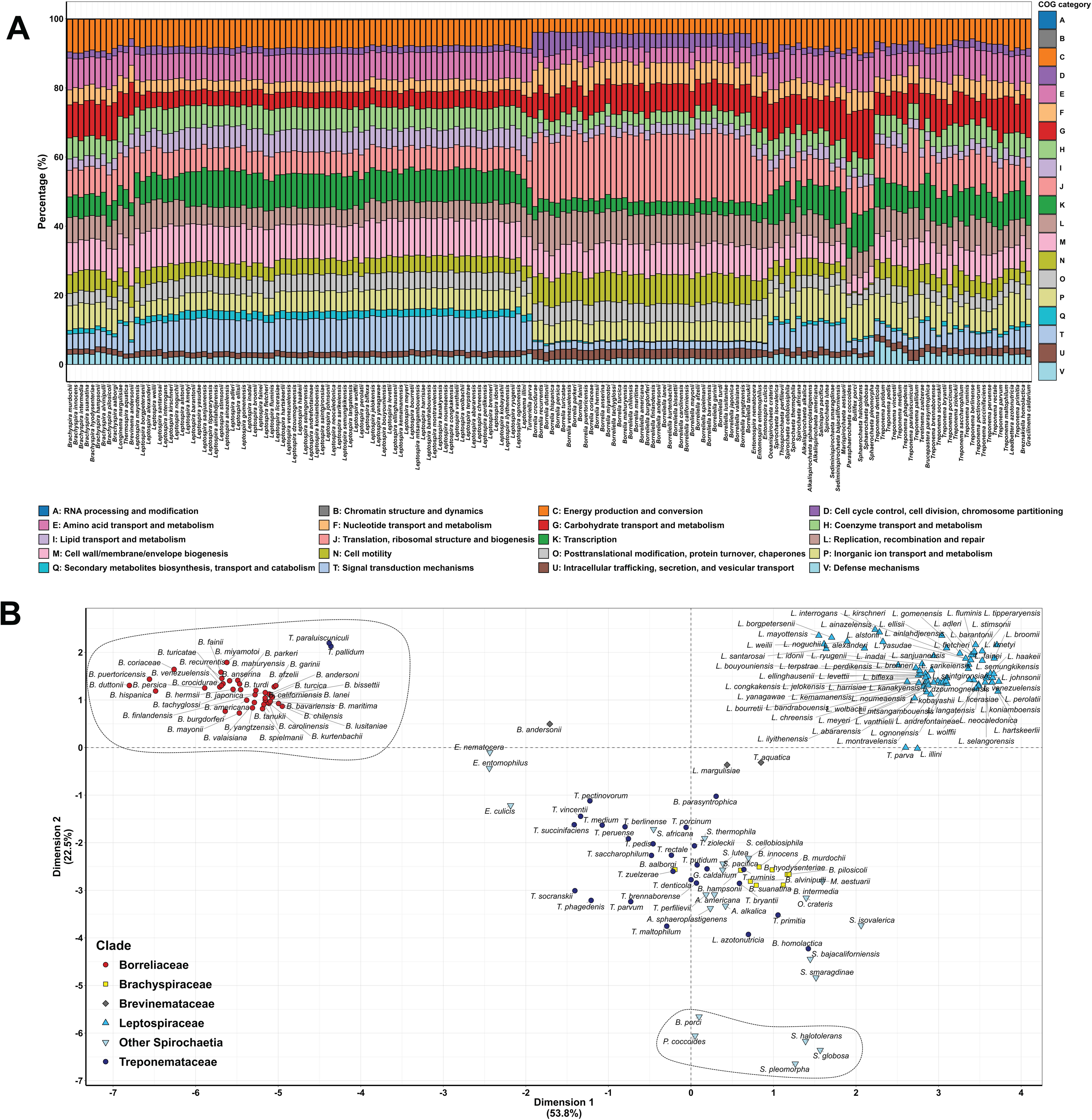
Functional genomic architecture differentiates ecological patterns. (A) Genome repartition by COG categories, excluding unknown function genes. The genomes are ordered from left to right according to their phylogenetic position in Fig 2, and COG categories are color-coded and ordered alphabetically from top to bottom as represented in the legend. The full name of each COG categories is presented at the bottom alongside their legend colours. This figure is also available on the website <u>spirochase.pasteur.cloud</u> for interactive exploration. (B) Principal Component Analysis (PCA) individuals plot of the COG category distribution, with the main taxonomic clades coloured as indicated in the legend. The clusters containing the host-dependent species (*Borrelia* spp., *T. pallidum* and *T. paraluiscuniculi*) (upper-left position) and the non-spiral *Spirochaetes* (lower position) are circled with a dashed line.

PCA analysis of individuals revealed clusters that only partially recapitulate phylogenetic relationships (Fig 3B). Instead, genomes tend to group according to key phenotypic and ecological traits. Host-dependent species, including *Treponema pallidum*, *Treponema paraluiscuniculi*, and members of the *Borreliaceae*, formed a distinct cluster despite their phylogenetic distance, consistent with convergent functional adaptation to parasitic lifestyles (Fig 3B). This clustering is not solely explained by genome reduction, as species with similarly small genomes (e.g. *Entomospira* spp. or *Brevinematales*) occupy distinct regions of the functional space (Fig 1A, Fig 3B). Similarly, all non-spiral *Spirochaetes* clustered together, independently of their phylogenetic position within *Spirochaetia*, indicating a shared functional signature associated with their unique shape phenotype in the phylum. *Leptospira* species also formed a separate cluster, reflecting their distinct functional repertoire (Fig 3B, Table S4). Altogether, these results show that functional genome composition captures phenotypic and ecological convergence across the phylum more effectively than phylogenetic relationships alone, particularly in lineages that have undergone strong ecological specialization.

### Functional genome allocation distinguishes host-dependent lifestyles in Spirochaetes

Once we identified functional signatures across the *Spirochaetes* phylum, we characterized them using enrichment analysis of functional categories. Here, enrichment is defined as the relative proportion of the genome allocated to a given functional category (COG), rather than the proportion of shared genes, allowing that different species may converge on similar functional profiles through different gene repertoires.

Host-dependent species showed a significant enrichment in categories related to translation (J) and cell cycle control (D), together with a depletion in secondary metabolism (Q), signal transduction (T) and transcription (K) (Fig 4A). In contrast, *Leptospiraceae* were enriched in secondary metabolism (Q) and chromatin dynamics (B) and displayed a reduced proportion of genes devoted to translation (J) or carbohydrate metabolism (G) (Fig 4B). This is consistent with previous studies showing that the *Leptospira* genus are the only spirochetes encoding for histone-like proteins involved in chromatin organization (38). Non-spiral *Spirochaetes* showed a marked depletion in motility functions (N, log_2_FC of -3.8), alongside an enrichment in carbohydrate transport and metabolism (G) (Fig 4C). Notably, motility-related genes were nearly absent in these genomes, indicating that the loss of the spiral morphology is tightly associated with loss of motility functions (Fig 4C, Table S4).

**Figure 4.**
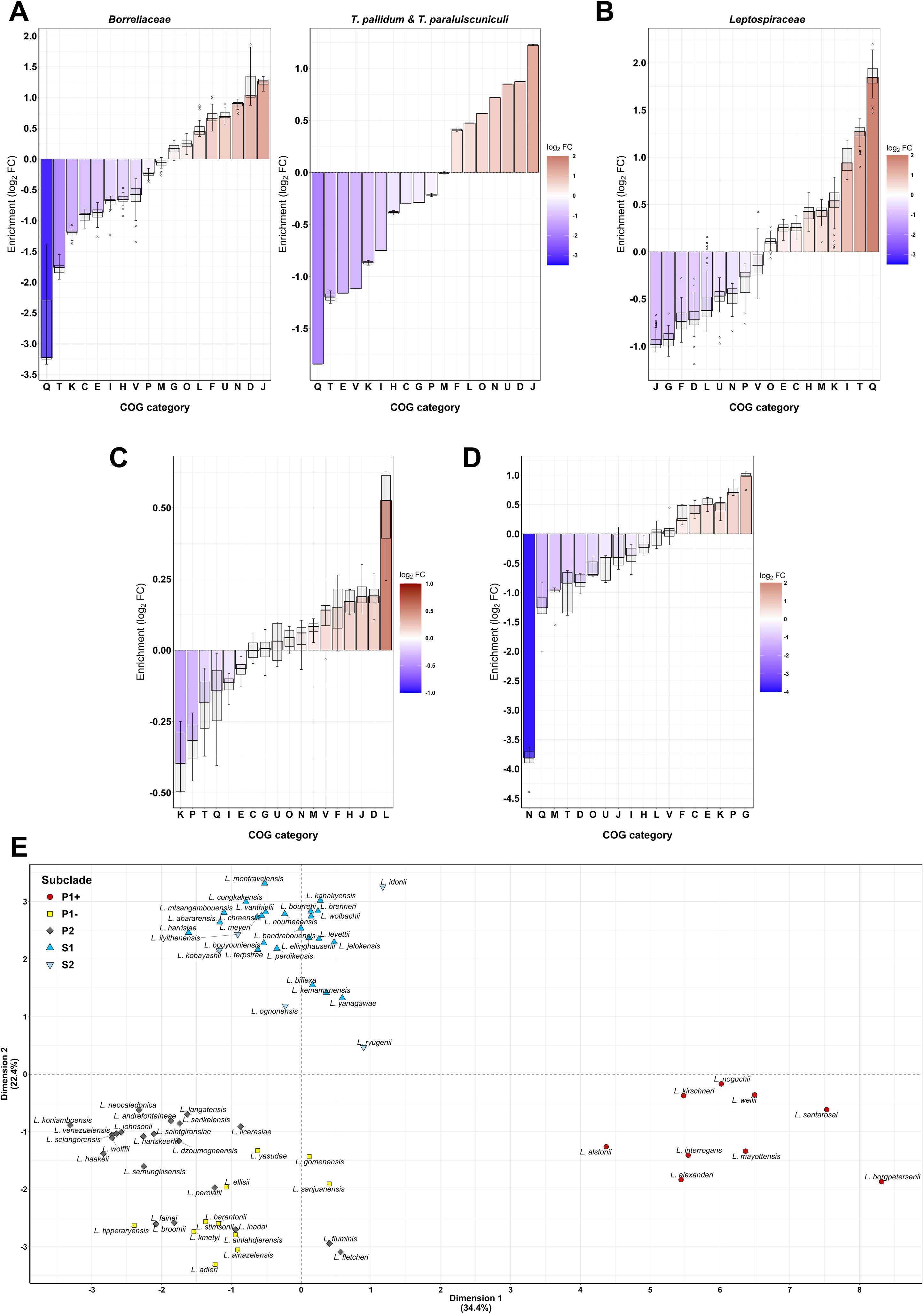
Functional genomic signatures of host-dependent phenotypic adaptation. (A) Enrichment analysis of the COG categories of the *Borreliaceae* family, *T. pallidum* and *T. paraluiscuniculi* (excluding *Borreliaceae* spp.) against all other *Spirochaetes* species. (B) Enrichment analysis of the COG categories of the *Leptospiraceae* family against all other *Spirochaetes* species. (C) Enrichment analysis of the COG categories in the non-spiral Spirochaetes (*B. porci*, *P. coccoides*, *S. pleomorpha*, *S. globosa* and *S. halotolerans*) versus all other *Spirochaetes*. (D) Enrichment analysis of the COG categories in the P1+ *Leptospira* spp. versus all other *Leptospira* spp. Log_2_FC values are arranged from lower to higher (left-to-right) and coloured according to the legend. The boxplots indicate the log_2_FC median and the first and third quartiles for all species within that group. Enrichment values (expressed as log_2_FC) are arranged by increasing order (left-to-right) and coloured according to the colour gradient on the left of each plot. The boxplots indicate the log_2_FC median and the first and third quartiles for all evaluated species. COG category letters correspond to: [A] RNA processing and modification, [B] Chromatin structure and dynamics, [C] Energy production and conversion, [D] Cell cycle control, cell division, chromosome partitioning, [E] Amino acid transport and metabolism, [F] Nucleotide transport and metabolism, [G] Carbohydrate transport and metabolism, [H] Coenzyme transport and metabolism, [I] Lipid transport and metabolism, [J] Translation, ribosomal structure and biogenesis, [K] Transcription, [L] Replication, recombination and repair, [M] Cell wall, membrane, envelope biogenesis, [N] Cell motility, [O] Posttranslational modification, protein turnover, chaperones, [P] Inorganic ion transport and metabolism, [Q] Secondary metabolites biosynthesis, transport and catabolism, [T] Signal transduction mechanisms, [U] Intracellular trafficking, secretion, and vesicular transport, and [V] Defense mechanisms. (E) Principal Component Analysis (PCA) individuals plot of the COG category distribution within *Leptospira* spp., with the main subclades coloured as represented in the legend.

To further illustrate the resolution of this functional classification, we applied the same approach within the genus *Leptospira*. To date, no study has previously been able to distinguish *a priori* highly virulent leptospires (P1+ group) from non-pathogenic species of the P clade (P1- group and P2 subclade) based solely on their genomic features. Here, COG categories distributions revealed clearly distinct functional signatures between these groups, with all P1+ *Leptospira* spp. clustering together, while P1- species grouped with P2 species despite belonging to a different phylogenetic subclade (Fig 2, Fig 4D, Fig 4E). Notably, *L. borgpetersenii* serovar Hardjo-Bovis, the most host-dependent *Leptospira* species (39), showed the most differential functional divergence from other pathogenic leptospires, suggesting that stronger ecological specialization is reflected in more pronounced functional genome allocation patterns (Fig 4E).

Overall, these results demonstrate that genome-wide functional allocation patterns, rather than gene content alone, provide a robust, sensitive and generalizable signature for ecological specialization and lifestyle diversification in *Spirochaetes*.

### Ancestral reconstruction and phylogenetic profiling uncover conserved spirochete traits

To further investigate the evolution of *Spirochaetes*, we used a conservative maximum-likelihood-based ancestral reconstruction approach to identify OGs present at the ancestral root of the phylum (LSCA) (Fig 2). Among the 34124 OGs analyzed, 511 OGs (1.5%) were inferred as ancestral to the whole phylum, with a probability above 80% for 60% of them (*N*=304) (Fig S12, Table S5). The most represented functions (31%, *N*=159) were housekeeping processes such as DNA replication (L), transcription (K) and translation (J) (Fig 5).

**Figure 5.**
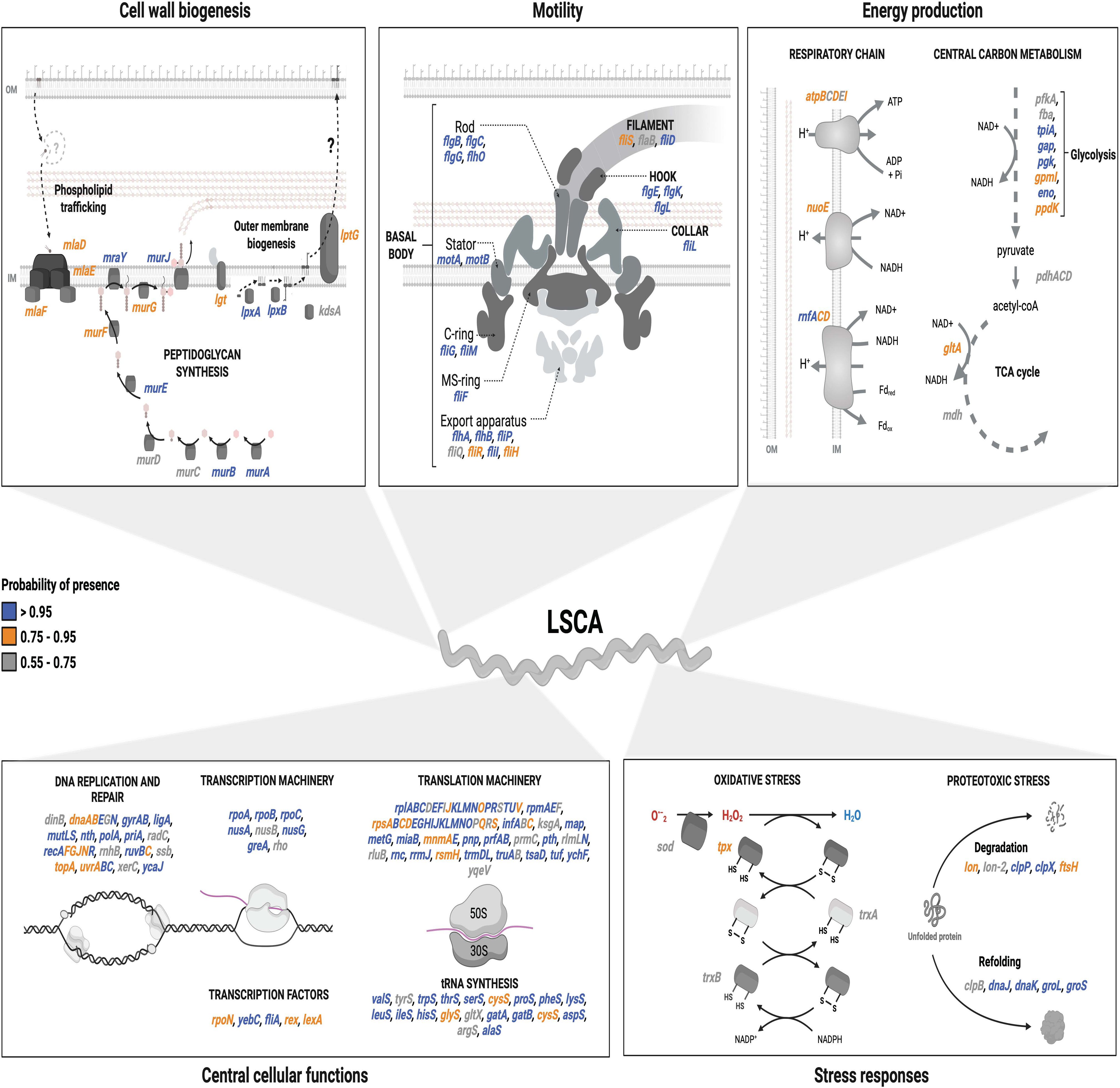
Features of the Last Spirochaetal Common Ancestor (LSCA). Schematic representation of the metabolism and selected features of the LSCA as inferred from the OGs present in the root. Each gene name is coloured according to their probability of presence indicated in the legend on the left-side. Each of the five panels represents a subset of genes with related functions. Question marks indicate missing functions classically needed for the pathways of cell wall biogenesis.

We also identified 23 ancestral flagellar proteins, representing a nearly complete flagellar apparatus and supporting a motile LSCA (Fig 5, Table S5). Notably, genes encoding the L- and P-ring components (FlgH and FlgI) were consistently absent from the reconstructed ancestral genome (Fig 5). This absence is unlikely to reflect limited methodological sensitivity, as most other flagellar components were recovered with high confidence (>95% for 78% of them, Table S5), and multiple orthologs of FlgH, FlgI and FlgJ were correctly annotated across all *Spirochaetes*. Specifically, we annotated 4 putative FlgH orthologs (OGs 11125, 12021, 12664 and 33851), 3 FlgI orthologs (Ogs 5854, 6973, and 12267), and 9 FlgJ orthologs (OGs 16615, 21301, 22375, 20264, 20665, 22998, 24731, 27829, and 31845). However, their inferred absence probabilities at the root ranged from 90-99% for FlgH, 57-96% for FlgI, and 80-98% for FlgJ, indicating very limited support for vertical inheritance from the LSCA in most cases, except for a single FlgI ortholog (OG12267; *N*=4 species, 57.6% absence probability). This pattern is also consistent with the revisited phylogenetic placement of early-branching lineages such as *Brachyspira* and *Brevinematales*, suggesting that the LSCA may have lacked flagellar structures traversing the peptidoglycan (Fig 2). Supporting this, domain annotation identified that all FlgJ orthologs lacked the muramidase domain required for peptidoglycan penetration in externally flagellated systems. Together, these observations are consistent with a periplasmic, endoflagellar configuration in the LSCA, although alternative evolutionary scenarios cannot be excluded, especially for components with weaker support (Fig 5, Table S5).

In addition to these ancestral functions, we recovered genes involved in cell wall biogenesis, including peptidoglycan production (*murABCDEFGJ*, *mraY*), enzymes of lipid A biosynthesis (*lpxAB*, *lgt*, *kdsA*, *htrB*), components of the Maintenance of Lipid Assymetry pathway (*mlaDEF*), and a homolog of the LPS permease LptG, most with probabilities above 80-90% (Fig 5). Cytoskeletal proteins associated with either membrane or cell curvature (*mreB, ccmA*) were also present with high probability (>98%), consistent with a putatively spiral-shaped ancestor (40,41). Genes involved in metabolic pathways, such as glycolysis, TCA cycle, and the respiratory chain, further indicate that the LSCA was a heterotrophic bacterium capable of central carbon metabolism (Fig 5). Finally, we identified genes encoding oxidative (*sod*, *tpx*, *trxAB*) and proteotoxic (*dnaKJ*, *groEL*, *clpB*) stress-response factors, together with two heme-related enzymes (*hemNH*). Together, these findings indicate that the LSCA combined a conserved housekeeping core with features consistent with an environmentally versatile, heterotrophic, motile, and likely spiral-shaped bacterium.

To complement this ancestral reconstruction, we generated a comprehensively annotated ortholog database that is made available to the community interested in spirochete research in the site <u>spirochase.pasteur.cloud</u>. This website allows the community to search for any ortholog of interest, explore its phylogenetic profile on the phylum, and extract its available functional information. To illustrate the combined versatility of these datasets, we used phylogenetic profiling to generate functional predictions over previously uncharacterized genes. We searched for genes conserved in most spirochetes but absent from the non-spiral lineages. As expected, most candidate genes identified by this approach were motility-related, confirming the internal consistency of the strategy.

Beyond these, several highly conserved gene families of unknown function also emerged from this analysis, representing interesting candidates for future experimental characterization (Fig S13, Table S6). These results illustrate how comparative functional genomics can generate testable hypotheses about conserved but functionally uncharacterized spirochete traits, with the full list of candidates available in Table S6 and through the <u>spirochase.pasteur.cloud</u> resource.

Overall, the analyses here presented demonstrate that integrating phylogenomic reconstruction with phylum-wide functional genome profiling provides a powerful framework for linking genome architecture to ecological transitions in bacteria.

## DISCUSSION

In this study, we showed that functional genome architecture provides a robust and generalizable lens for interpreting ecological diversification in bacteria, a finding demonstrated here across the full ecological breadth of the very diverse *Spirochaetes* phylum. We determined that the proportional allocation of coding capacities across functional biological processes helps predict ecological lifestyles and phenotypic diversification in *Spirochaetes* independently of phylogenetic relationships.

Despite syphilis, Lyme disease and leptospirosis being emerging or re-emerging diseases, little is known about the biology of spirochetes compared to model bacteria. Spirochete research in both pathogenic and non-pathogenic species has traditionally been hampered by fastidious growth, limited genetic tools and incomplete genome annotations, complicating the progress of hypothesis-driven studies following omics analyses (42,43). To address this, we significantly improved the functional annotation of spirochaetal proteins by applying a cluster-based approach, achieving an average 70% annotation level per species and nearly 90% annotation for some genomes. This represents a significant improvement (∼15%) over previous studies, where, for example, only 44% of the *L. interrogans* genome was annotated, compared to 64% in our analysis (24,44). In addition, we assigned putative origin (bacterial, eukaryotic, viral or archaeal) and closest ortholog for 60% (*N*=8496) of genes of unknown function (S), reducing the proportion of orphan genes in *Spirochaetes* to just 16% (*N*=5735), a significant advancement for the field.

The improved annotation coverage enabled the detection of potential lineage-specific genomic acquisitions with ecological significance. *Leptospira* spp. showed a significantly higher number of ortholog groups of eukaryotic origin compared to other *Spirochaetes*, including proteins associated with actin cytoskeleton dynamics or growth regulation, functions typically absent or rare in bacterial genomes. While this pattern may partly reflect annotation bias, whereby divergent bacterial sequences are preferentially assigned to eukaryotic homologs due to database composition, the consistent enrichment observed in all *Leptospira* spp. also supports the presence of *bona fide* eukaryotic-like genes. Although the frequency and directionality of inter-kingdom gene transfers is often difficult to resolve, horizontal gene exchange between eukaryotes and bacteria has been documented in several host-associated microorganisms (35,45), raising the possibility that such events of acquisition contributed to the evolution of this lineage. *Leptospira* spp. were previously estimated to appear as environmental bacteria (46). Therefore, they may have exchanged genetic material over long periods in the soils, which harbour a remarkably diverse array of eukaryotes, ranging from plants and fungi to protists and animals, encompassing multiple major lineages across the eukaryotic tree. Pathogenic leptospires subsequently emerged with the rise of mammals and, over long periods, adapted in some cases to animal reservoirs. Their co-evolution with eukaryotes over hundreds of millions of years may be reflected here in this enrichment of eukaryotic-like proteins. Although the ecological context differs, in *Legionella* species, horizontal gene transfer with their natural hosts (free-living protozoa) has shaped their genomes, which contain a high number of genes encoding proteins with eukaryotic domains, reflecting the long-term co-evolution of these microorganisms (47). Whether these eukaryotic acquisitions in *Leptospira* spp. were retained through selective advantage in host interaction, niche adaptation, or modulation of eukaryotic cellular processes remains to be determined, but they may represent yet another distinctive feature of *Leptospira* genome evolution relative to the rest of the phylum (36). Conversely, *Borreliaceae* showed a significant enrichment in ortholog groups of viral origin, consistent with the abundance of phage-derived plasmids in these species (21). These elements are known to contribute to genome plasticity and have been implicated in host adaptation and immune evasion in *Borrelia* species (48–50), suggesting that viral genetic material has been repeatedly selected as an adaptive resource during the evolution of this lineage. Together, these findings indicate that horizontal acquisition of foreign genetic material has contributed to shaping the distinct genomic identities of these lineages.

In this study we provide a robust comprehensive and updated phylogeny for the *Spirochaetes* phylum supported by multiple comparative genomic methods. This revisited phylogeny redefines the evolutionary rooting of the phylum relative to previous studies, likely reflecting the increased species coverage and more adequate selection of the outgroup (Fig 2) (2), and provides a basis for revising its taxonomy. We propose a 45% POCP threshold for genus delineation within *Spirochaetes*, which would resolve a long-standing controversy over genus classification (51–54). Applying this threshold consistently reveals that the *Treponema* genus, as currently defined, aggregates lineages with substantially different levels of genomic divergence, while conversely supporting the unification of currently separated taxa within *Borreliaceae*. We therefore strongly recommend either standardizing genus nomenclature across the phylum under objective genomic criteria or, alternatively, explicitly recognizing that the rank “genus” within *Spirochaetes* reflects varying levels of species divergence depending on the lineage. For example, we show that the *Leptospira* genus encompasses greater diversity than the *Borreliaceae* family (which includes the genera *Borrelia* and *Borreliella*).

These inconsistencies highlighted the limits of phylogenetic and genomic similarity metrics alone in capturing the observed functional and ecological diversification within *Spirochaetes*. Here, we demonstrate that functional genome composition correlates with ecological lifestyles, revealing convergent functional adaptation between distant lineages, consistent with the response to parallel selective pressures. Indeed, we show that genome-wide functional allocation patterns, defined as the proportional coding capacity devoted to each functional category, correlate with ecological lifestyle and phenotypic traits across *Spirochaetes*, independently of gene content or phylogenetic relatedness. Host-dependent lineages converge on a shared functional profile characterized by enrichment in translation and cell cycle control categories and depletion in transcription and secondary metabolism, despite their phylogenetic distance. The simplest interpretation is that adaptation to obligate host dependence imposes common constraints on coding functional allocation, regardless of the ancestral genomic background. Consistently, we showed that host-dependent species allocate a larger fraction of their genome to translation and a reduced one to transcription. This may suggest that parasitic microorganisms have shifted genome regulation towards reducing transcription factors but increasing post-transcriptional regulation, as evoked by *Borrelia* spp. (48,55–58).

Importantly, this functional framework also revealed finer ecological differences within the genus *Leptospira* that had previously resisted genomic classification. Highly virulent P1+ leptospires display a distinct functional profile from non-pathogenic species of the P1 clade. This distinction emerges from functional genome allocation alone, without recourse to phenotypic data, indicating that ecological specialization leaves a measurable imprint on the genomic architecture of P1+ species that existing classification approaches had not previously captured (46). Additionally, the most host-dependent *Leptospira* species, *L. borgpetersenii* serovar Hardjo-Bovis, shows the most pronounced functional divergence from other pathogenic leptospires. This relationship between degree of host dependence and magnitude of functional divergence suggests that functional genome remodeling is not a binary transition but a gradual process that evolves as ecological constraints become more restrictive. Whether these functional signatures reflect a universal principle of microbial specialization remains to be tested, but their predictive power is demonstrated here across the ecological diversity of a phylum.

Beyond these evolutionary insights, the annotated ortholog database generated here provides a practical resource for characterizing previously uncharacterized gene families in *Spirochaetes*. This resource, as applied here by comparing gene conservation patterns between spiral and non-spiral lineages, offers a systematic route to identifying candidate functions among the large proportion of conserved hypothetical proteins that remain uncharacterized in this phylum. Whether any of the candidate families identified through this approach contribute to motility, cell shape, or other spirochete-specific traits remains to be determined experimentally, but the strategy demonstrates the predictive value of integrating functional and phylogenetic information at the phylum scale. The full annotated ortholog database is available at spirochase.pasteur.cloud.

Lastly, reconstruction of the genome of the Last Spirochaetal Common Ancestor revealed a motile, diderm, heterotrophic bacterium with a nearly complete flagellar apparatus, a conserved housekeeping core, and stress response systems consistent with its emergence around the Great Oxidation Event (∼2300 MYA) (59). The LSCA possessed a SOD-Trx system for oxidative stress resistance and two heme-related enzymes (*hemNH*), suggesting a partial heme biosynthesis capacity subsequently lost in modern *Borrelia* and *Treponema* (60). The absence of L- and P-ring flagellar components from the reconstructed ancestral genome, supported by the lack of a muramidase domain across all identified FlgJ orthologs and consistent with the phylogenetic placement of early-branching lineages (14), is coherent with a periplasmic endoflagellar configuration in the LSCA, and aligns with most updated models on flagellar evolution (61,62). Of note, while cytoskeletal proteins such as MreB and bactofilins were identified in the LSCA, their presence alone is insufficient to conclude a spiral morphology, as these proteins influence membrane curvature but do not universally determine cell shape across bacteria. Indeed, *B. burgdorferi* encodes bactofilins yet its spiral shape depends primarily on flagellar mechanics (63). The determination of the spiral shape of spirochetes is therefore likely multifactorial and genus-dependent, involving both flagellar and peptidoglycan-associated proteins (41,64,65), some of which may correspond to the gene families identified here as absent in non-spiral lineages, representing interesting candidates for future investigation. Regardless of the mechanism, the monophyly of all non-spiral species and their restriction to a single clade within *Spirochaetia* provides sufficient evidence to conclude that the spiral morphology is ancestral to the phylum and was lost only once (Fig 2). The presence of only a partial LPS biosynthesis and export pathway in the LSCA is insufficient to conclude on LPS presence, consistent with *Leptospira* being the only known LPS-containing *Spirochaetes* (66). Together, these inferred ancestral features shed light on the ecology and lifestyle of the earliest spirochetes, and on the evolutionary trajectories of some of the hallmark traits of modern spirochetes.

The present framework, built on cultivable representatives with high-quality genomes, provides a robust reference for future integration of metagenome-assembled genomes from uncultivated spirochete lineages, which may further refine the evolutionary picture presented here and could reveal additional functional transitions associated with unexplored ecological niches in *Spirochaetes*. Overall, this study provides a reference framework for spirochete research while illustrating how functional genome composition can be used to track ecological diversification across the breadth of a bacterial phylum.

## METHODS

### Data collection

176 genomes covering all cultivable species within the *Spirochaetes* phylum were downloaded from the National Center for Biotechnology Information (NCBI) as of August 2024 following manual curation. The genomes corresponding to the species *Alkalispirochaeta odontotermitis* (GCF_000768055.1), *Rectinema subterraneum* (GCF_009768935.1), *Treponema lecithinolyticum* (GCF_000468055.1), and *Marispirochaeta associata* (GCA_001749745.1) were removed because of high level contamination with other organisms. For the selection of genomes, one genome was chosen per species TaxID in NCBI prioritizing the RefSeq-validated genome per TaxID. For the two species in which there was no RefSeq-validated reference genome (*Borrreliella chilensis* and *Longinema margulisiae*), we inspected the literature and selected the only available GenBank genome after checking completeness and contamination. For the reference genomes of the 69 *Leptospira* spp., we followed the latest Position Statement of the International Leptospirosis Society (ILS) (67) and the International Committee on Systematics of Prokaryotes Subcommittee on the taxonomy of *Leptospiraceae* (68). It should be noted that there is no publicly available genome for the genus *Cristispira*. The final selected number of genomes is 172 and their description and assembly numbers are available in Table S1.

### Orthology inference

For the orthology inference, we employed the protocol described in (69) with a few modifications. We used OMA standalone version 2.5.0 (25) with the exported reference proteomes. To ensure that orthologs were computed without bias, in a first run we used only the 172 pre-selected proteomes with the following parameters: estimated species tree, bottom-up inference of HOGs, mid-point rooting, minimum score of 181, length tolerance ratio of 0.61, stable pair tolerance of 1.81, inparalog tolerance of 3.0, verified pair tolerance of 1.53, minimum sequence length of 50, and minimum edge completeness of 0.65. In a second run, we used the 172 pre-selected proteomes along with two *Lindowbacteria* species (GCA_001782795, GCA_001784175) used as outgroup for rooting, with the same parameters as above except for the rooting. For the orthology inference of *Treponematales*, we used the 28 pre-selected genomes belonging to the *Treponemataceae* clade (Table S1) alongside three of their closest relatives for the rooting (GCF_000143985.1, GCF_000378205.1, GCF_002087085.1) (Fig 2) with the same parameters as above.

Calculation of the pangenome accumulation curve was obtained following the method described in (70), using 100 random iterations in the presence/absence matrix of OGs. The regression curve was calculated using a generalized additive model (gam) with a cubic spline under the formula y ∼ s(x, bs = “cs”).

### Tree inference

Tree inference was determined as described in (69) with a few modifications. Briefly, we first selected 140 highly-conserved soft-core genome markers (95% conservation, 163 species) from the first run of OMA OGs using the filter_groups.py script from (69) and aligned them individually with MAFFT version 7.505 under the L-INS-i algorithm (71). We then soft-trimmed the alignments individually using BMGE version 1.12 (72) under the BLOSUM30 matrix with the following parameters: maximum entropy threshold of 0.95, sliding window size of 1, minimum length of selected regions of 1. Lastly, we concatenated the trimmed alignments to build the supermatrix using the concat_alignments.py script available in (69). This supermatrix was then used as input for IQ-TREE version 2.3.2 (73) with 10,000 ultra-fast bootstraps and 10,000 SH-alrt (Shimodaira-Hasegawa-like procedure test) (74) under the homologous best-fit bacterial model of evolution LG+F+I+R10. The tree generated was then used as a guide tree for a second tree inference under the heterogeneous model of evolution LG+C20+R10 with IQ-TREE and the same parameters. The same process was repeated with the second run of OMA OGs including the two *Lindowbacteria* genomes (see above) with 130 highly conserved soft-core genome markers (95% conservation, 165 species).

For the coalescence-based method, we used the same 130 trimmed soft-core genome marker alignments including the two *Lindowbacteria* genomes (see above) and inferred individual phylogenies using FastTree version 2.1.11 (75). We then used these individual phylogenies to infer the final species tree using ASTRAL version 5.7.8 (76).

For the *Treponematales* phylogenetic inference, we used 440 soft-core genome markers (95% conservation, 29 species) obtained from the orthology inference (see above) and proceeded as before to obtain the supermatrix. This supermatrix was then used as input for IQ-TREE version 2.3.2 with 10000 ultra-fast bootstraps and 10000 SH-alrt (Shimodaira-Hasegawa-like procedure test) under the homologous best-fit bacterial model of evolution LG+F+I+R10.

To support the phylogenetic tree, we calculated pairwise comparisons of Percentage of Conserved Proteins (POCP), Average Nucleotide Identity (ANI) and a kmer-based method (TETRA). ANI (with the BLASTn module) and TETRA were calculated with PyAni version 0.2.11 (77). POCP was calculated with POCP-nf version 2.2.0 (32,78).

### Functional annotation

Functional annotation of the proteomes was determined using a cluster-based approach with fine-grained orthology. This strategy assumes that all sequences within an ortholog group share a common function, allowing functional information available for a single well-characterized sequence to be propagated across all members of the OG. This is particularly advantageous for divergent proteins that may not reach the identity thresholds required for direct database hits in classical annotation pipelines. However, this approach requires ortholog groups inferred with high precision (that is, with a high fraction of true orthologs and minimal paralog contamination). OMA-based ortholog groups are therefore well-suited for this purpose, given their stringent evolutionary-aware inference and low false-positive rates. Combined with the fine-grained ortholog annotations derived from eggNOG and eggNOG-mapper, this strategy improves annotation quality and coverage compared to other sequence-similarity-only pipelines.

Briefly, we obtained a single representative sequence per ortholog group by clustering each OG from the OMA orthology inference (see above) using MMseqs2 version 15-6f452 (79) with the following parameters: workflow easy-cluster, infinite e-value, minimum sequence identity 0, coverage mode 1, alignment coverage 0. Each representative sequence was then searched with eggNOG-mapper version 2.1.12 (80) on eggNOG DB version 5.0.2 (81). Sequence searches were performed using four consecutive approaches: DIAMOND ultra-sensitive (82), HMMER with database 2 (bacteria), HMMER with database 2157 (archaea), and HMMER with database 10239 (viruses) (83). We then assigned an annotation to each OG according to the following hierarchy: if the DIAMOND search provided a COG category different from “unknown” (S/NA) it was retained, otherwise we looked for the unannotated OGs in the HMMER searches (first bacterial, then archaeal, lastly viruses). The DIAMOND search provided an annotation for 53.5% of all OGs, and the HMMER searches complemented this to 58.2% of all OGs. For assignment of the maximum annotation level following eggNOG-mapper, the same hierarchical procedure was followed.

Principal Component Analysis (PCA) was performed with the PCA function of the FactoMineR package (84) in R version 4.3.0 (85) with the normalized percentage of known COG categories per genome as input. Log_2_FC values were calculated as the logarithmic base 2 ratio between the percentage of OGs within a species’ genome annotated as a given COG category and the percentage of OGs within all other species’ genomes annotated as the same COG category, unless otherwise stated.

### Ancestral genome reconstruction

The genomic components present in the LSCA were reconstructed using PastML version 1.9.41 (86) with a combination of the presence/absence matrix of OGs and the phylogeny presented in Fig 2. To ensure a robust inference, ancestry prediction was performed in two steps: first, we run PastML with a marginal posterior probabilities approximation (MPPA method) to determine the maximum-likelihood model parameters and marginal probabilities for each column under the F81 model (87); then, we run PastML again using these pre-determined model parameters to evaluate the most likely evolutionary scenario and produce an internally consistent result for all nodes under the JOINT method (88). Therefore, the final root status is the one obtained under the JOINT prediction and the probabilities presented are the ones obtained under the MPPA prediction. For the final analysis, only OGs with JOINT-based root status “presence” and an MPPA-based probability ≥55% were retained.

### Phylogenetic profiling

To obtain the OGs lost by the non-spiral *Spirochaetes* (*Bullifex porci*, *Parasphaerochaeta coccoides*, *Sphaerochaeta pleomorpha*, *Sphaerochaeta globosa* and *Sphaerochaeta halotolerans*), we first looked for OGs that were completely absent from those species but present in 80% or more of the other *Spirochaetes* phylum species (Type = “ABSENT”). As a complementary strategy, we also looked for OGs completely absent from the same species but highly conserved (≥95% conservation) in the closest clade species (all *Treponemataceae* spp., *Alkalispirochaeta alkalica*, *Alkalispirochaeta americana*, *Alkalispirochaeta sphaeroplastigenens*, *Marispirochaeta aestuarii*, *Salinispira pacifica*, *Sediminispirochaeta bajacaliforniensis*, *Sediminispirochaeta smaragdinae*, *Spirochaeta africana*, *Spirochaeta lutea*, and *Spirochaeta thermophila*) (Type = “LOST”). Both datasets were then combined and are available in Table S6.

## AKNOWLEDGEMENTS

We thank Linda Grillova and Zachary Ardern for their support and valuable feedback through the earliest stages of development of this project. We thank the Biology of Spirochetes Unit at Institut Pasteur for their assistance in testing the *Spirochase* website and for their constructive feedback on the manuscript. We are grateful to Dr. Kevin Waldron and Dr. Pierre Garcia for helpful discussion and critically reading this manuscript. In the *Spirochase* website, RT oversaw UI/UX design and produced the interface mock-ups, and EC oversaw website implementation, development and deployment. We thank David Hampson and Kittitat Lugsomya for their help with finding new *Brachyspira* strains. This work was also supported by a PTR (Programmes Transversaux de Recherche) grant (PTR2019-310, NB) from Institut Pasteur (Paris, France), and by National Institutes of Health grant P01 AI-168148 (MP & NB). SGH is a recipient of the Pasteur-Paris University PhD program. KC has received a PhD scholarship from BioSPC doctoral school (ED562, Université Paris Cité). The authors acknowledge the use of AI-assisted copy-editing tools (LLMs) to support language editing, readability, formatting and style. All scientific content was developed and verified by the authors.

## SUPPLEMENTARY FIGURES

**Figure S1.** Pangenome analysis of the phylum *Spirochaetes*. (A) Pangenome accumulation plot of the *Spirochaetes* phylum representing the cumulative number of different OGs. This was calculated using 100 random iterations in the presence/absence matrix of OGs. Each blue dot represents one iteration, and the black line is the smooth curve of regression calculated using a generalized additive model (gam) with a cubic spline under the formula y ∼ s(x, bs = “cs”). (B) Cumulative (green dots) and non-cumulative (blue dots) numbers of orthologs shared as the number of species increases in the range 2 to 172. The Y axis is represented in logarithmic scale to facilitate visualization.

**Figure S2.** Phylogenetic comparisons of the *Spirochaetes* phylum (I). (A) Co-phylo plot representing the comparison between the phylogeny obtained under the unrooted homogeneous model of evolution (LG+F+I+R10, left side) and the unrooted heterogeneous model of evolution (LG+C20+R10, right side). Red lines connect the same leaves (species) in both trees. (B) Co-phylo plot representing the comparison between the phylogeny obtained under the rooted homogeneous model of evolution (LG+F+I+R10, left side) and the rooted heterogeneous model of evolution (LG+C20+R10, right side). Red lines connect the same leaves (species) in both trees.

**Figure S3.** Phylogenetic comparisons of the *Spirochaetes* phylum (II). (A) Co-phylo plot representing the comparison between the phylogeny obtained under the rooted homogeneous model of evolution (LG+F+I+R10, left side) and the unrooted homogeneous model of evolution (LG+F+I+R10, right side). Red lines connect the same leaves (species) in both trees. (B) Co-phylo plot representing the comparison between the phylogeny obtained under the rooted heterogeneous model of evolution (LG+C20+R10, left side) and the unrooted heterogeneous model of evolution (LG+C20+R10, right side). Red lines connect the same leaves (species) in both trees.

**Figure S4.** Phylogenetic comparisons of the *Spirochaetes* phylum (III). (A) Co-phylo plot representing the comparison between the phylogeny obtained under the rooted heterogeneous model of evolution (LG+C20+R10, left side) and the rooted multi-species coalescence model of evolution (ASTRAL, right side). Red lines connect the same leaves (species) in both trees. (B) Presence/absence plot of the 140 soft-core genome markers (OGs) used for the phylogenetic inference, with species in which the OG is present marked in blue and absent in grey. The Y axis contains the species ordered from top to bottom according to their phylogenetic position in Fig 2.

**Figure S5.** An updated phylogeny for the *Treponematales* order. Phylogeny of the *Treponematales* order obtained under the model LG+F+I+R10 (best-fit bacterial model for *Spirochaetes*) using 440 soft-core genome markers and rooted using the closest clade in Fig 2 (*Sediminispirochaeta smaragdinae*, *Sediminispirochaeta bajacaliforniensis*, *Marispirochaeta aestuarii*). The two subclades within the *Treponema* genus are highlighted in blue (T1 subclade) and grey (T2 subclade). All nodes had SH-alrt and bootstrap support values of 100% except for the one splitting *T. parvum* from *T. socranskii*, *T. porcinum* and *T. bryantii*, which had 97/91% support.

**Figure S6.** Average nucleotide identity matrix of the *Spirochaetes*. Phylogenetic tree of the cultivable species of *Spirochaetes*, with the main clades highlighted in colours (yellow for *Brachyspira* spp., dark grey for *Brevinematales*, cyan for *Leptospirales*, red for *Borreliaceae*, green for *Entomospira* spp., light blue for intermediate *Spirochaetia*, and dark blue for *Treponemataceae*), represented next to the pairwise matrix of Average Nucleotide Identity using the BLASTn method (ANIb, %). The colours in the ANIb matrix correspond to the continuous scale represented in the right. The intermediate coloured column represents the main clade classification.

**Figure S7.** Kmer-based tetranucleotide matrix of the *Spirochaetes.* Phylogenetic tree of the cultivable species of *Spirochaetes*, with the main clades highlighted in colours (yellow for *Brachyspira* spp., dark grey for *Brevinematales*, cyan for *Leptospirales*, red for *Borreliaceae*, green for *Entomospira* spp., light blue for intermediate *Spirochaetia*, and dark blue for *Treponemataceae*), represented next to the pairwise matrix of a kmer-based tetranucleotide method (TETRA, %). The colours in the TETRA matrix correspond to the continuous scale represented in the right. The intermediate coloured column represents the main clade classification.

**Figure S8.** Percentage of conserved proteins as predictor of genus boundaries in Spirochaetes. (A) Histogram of distribution of Average Nucleotide Identity using the BLASTn method (ANIb, %) values for all pairwise comparisons of the phylum *Spirochaetes*. The distribution of the values is a single mode skewed uniformly to the right. (B) Histogram of distribution of Percentage of Conserved Proteins (POCP, %) values for all pairwise comparisons of the phylum *Spirochaetes*. The values are distributed in three main modes with clear boundaries between them. (C) Histogram of distribution of kmer-based tetranucleotide method (TETRA, %) values for all pairwise comparisons of the phylum *Spirochaetes*. The values are distributed in two main modes, with the first skewed to the right and with no clear boundaries between them. (D) POCP matrix of all pairwise comparisons of the phylum *Spirochaetes* that include a POCP value ≥ 45%. Values lower than the 45% threshold are coloured in white, all the other colours in the POCP matrix correspond to the continuous scale represented in the right.

**Figure S9.** Annotation graph of all *Spirochaetes* species. (A) Annotation graph of genome repartition by COG categories, including unknown function genes (COG category S). The genomes are ordered from left to right according to their phylogenetic position in Fig 2, and COG categories are color-coded and ordered alphabetically as represented in the legend on the right. (B) Representation of the maximum annotation level repartition per genome, including unannotated genes (category S). The legend on the right illustrates the putative origin of the OGs per genome: B for bacterial, A for archaeal, V for viral and E for eukaryotic. The genomes are ordered from left to right according to their phylogenetic position in Fig 2.

**Figure S10.** Comparative analysis of the non-bacterial OGs in the main genera of *Spirochaetes*. Percentage of OGs of putative eukaryotic (panel A), archaeal (panel B) or viral (panel C) origin in *Borreliaceae* spp. (grey dots), *Leptospira* spp. (blue dots), *Treponema* spp. (green dots) or *Brachyspira* spp. (purple dots). **\*** p-value < 0.05, **\*\*** p-value < 0.01, **\*\*\*** p-value < 0.001, **\*\*\*\*** p-value < 0.0001. The data was analysed using a Brown-Forsythe and Welch ANOVA test with Dunnett’s T3 post-comparison test.

**Figure S11.** Contribution of the individual variables to the Principal Component Analysis (PCA). Percentage of contribution of each COG category to the PC1 and PC2 (panel A), PC1 only (panel B) and PC2 only (panel C). The red dashed line indicates the percentage of contribution if all variables contributed the same to the components (5%).

**Figure S12.** OGs present in the Last Spirochaetal Common Ancestor. Presence/absence plot of the 511 OGs found to be present in the LSCA, with species in which the OG is present or absent marked in blue and grey, respectively. The Y axis contains the species ordered from top to bottom according to their phylogenetic position in Fig 2. OGs are ordered by COG category and from most to least abundant. The colour-coded lower panel indicates the COG category according to the legend on the right.

**Figure S13.** OGs lost by non-spiral *Spirochaetes*. Distribution of the number of OGs found to be lost by the non-spiral *Spirochaetes* (*B. porci*, *P. coccoides*, *S. pleomorpha*, *S. globosa* and *S. halotolerans*) through the two approaches employed in the study (see Methods). OGs in the LOST type (95% conservation or higher in the closest clade spiral species) are coloured in dark blue while OGs in the ABSENT type (80% or higher conservation in all other spiral *Spirochaetes*) are coloured in light blue.

## SUPPLEMENTARY TABLES

**Table S1.** Genomes used in this study and their main features.

**Table S2.** Percentage of conserved proteins (POCP) across all *Spirochaetes* species.

**Table S3.** Annotation of the OGs of putative eukaryotic origin found in *Leptospira* spp.

**Table S4.** Relative percentage of the genome of each *Spirochaetes* species devoted to each COG functional category.

**Table S5.** Annotation of the OGs found present in the LSCA.

**Table S6.** OGs lost by the non-spiral *Spirochaetes*.

