## Supplementary figures and images for "Genomic functional architecture predicts ecological adaptation across a highly diverse bacterial phylum"

### FigS1

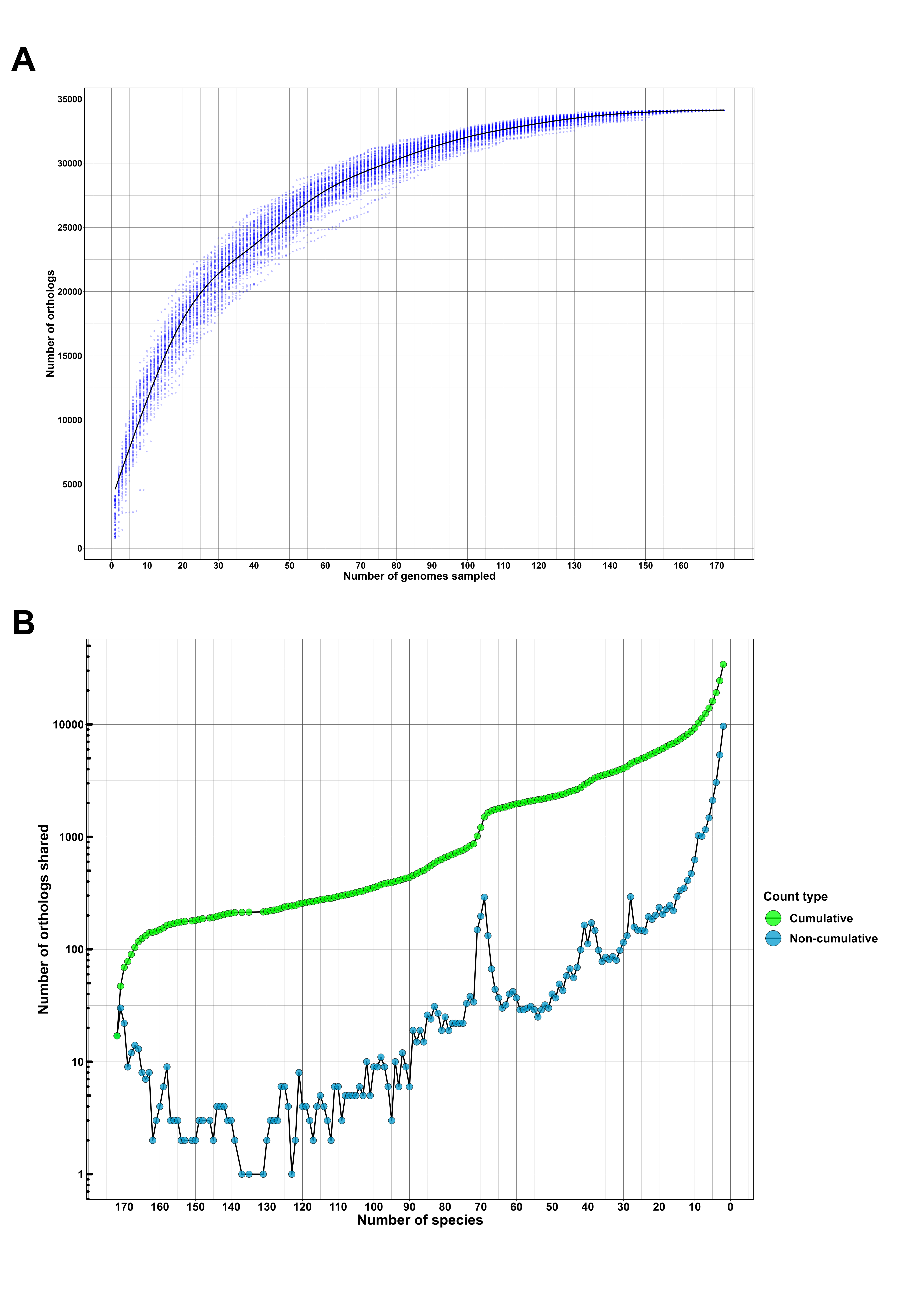

### FigS2

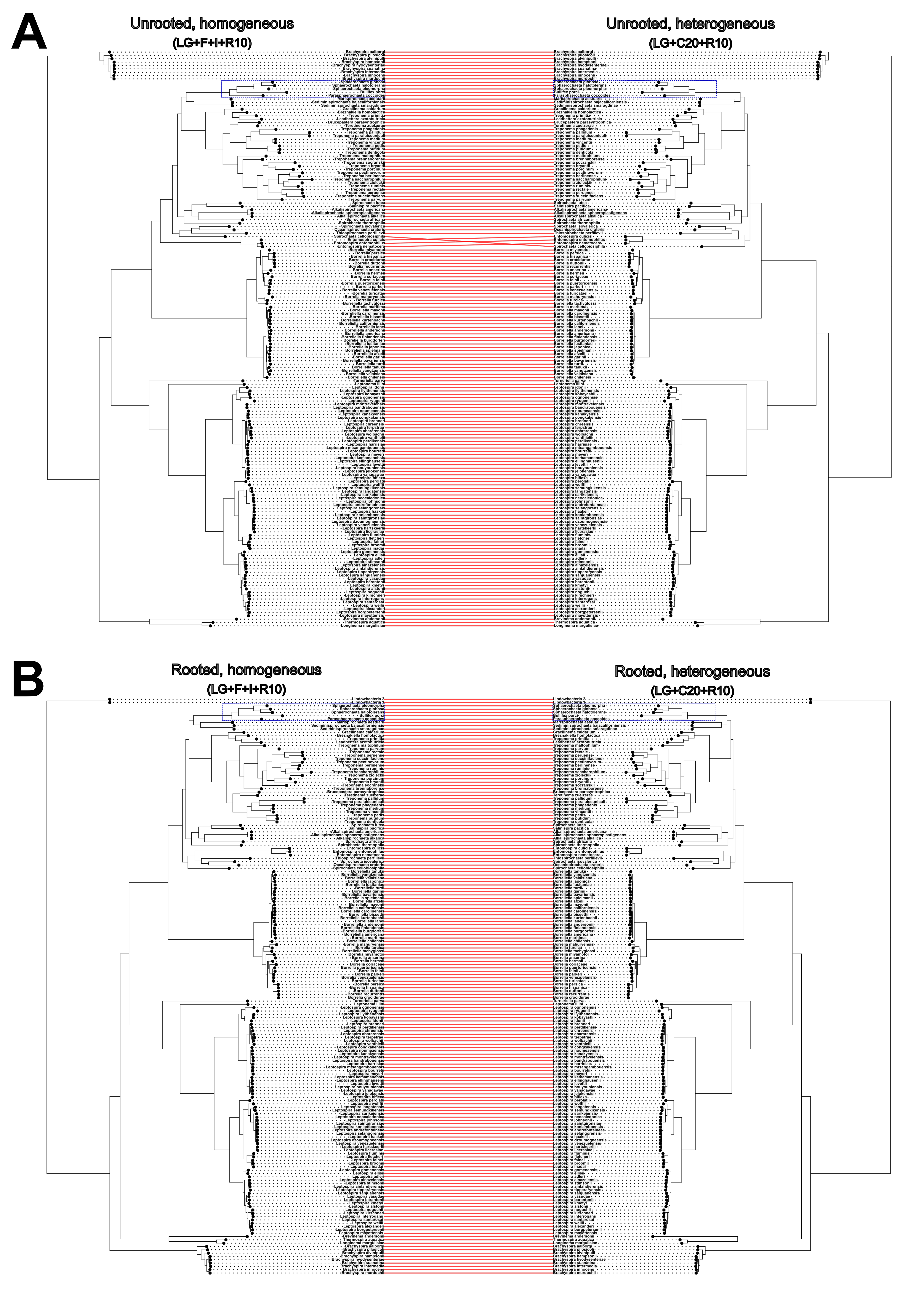

### FigS3

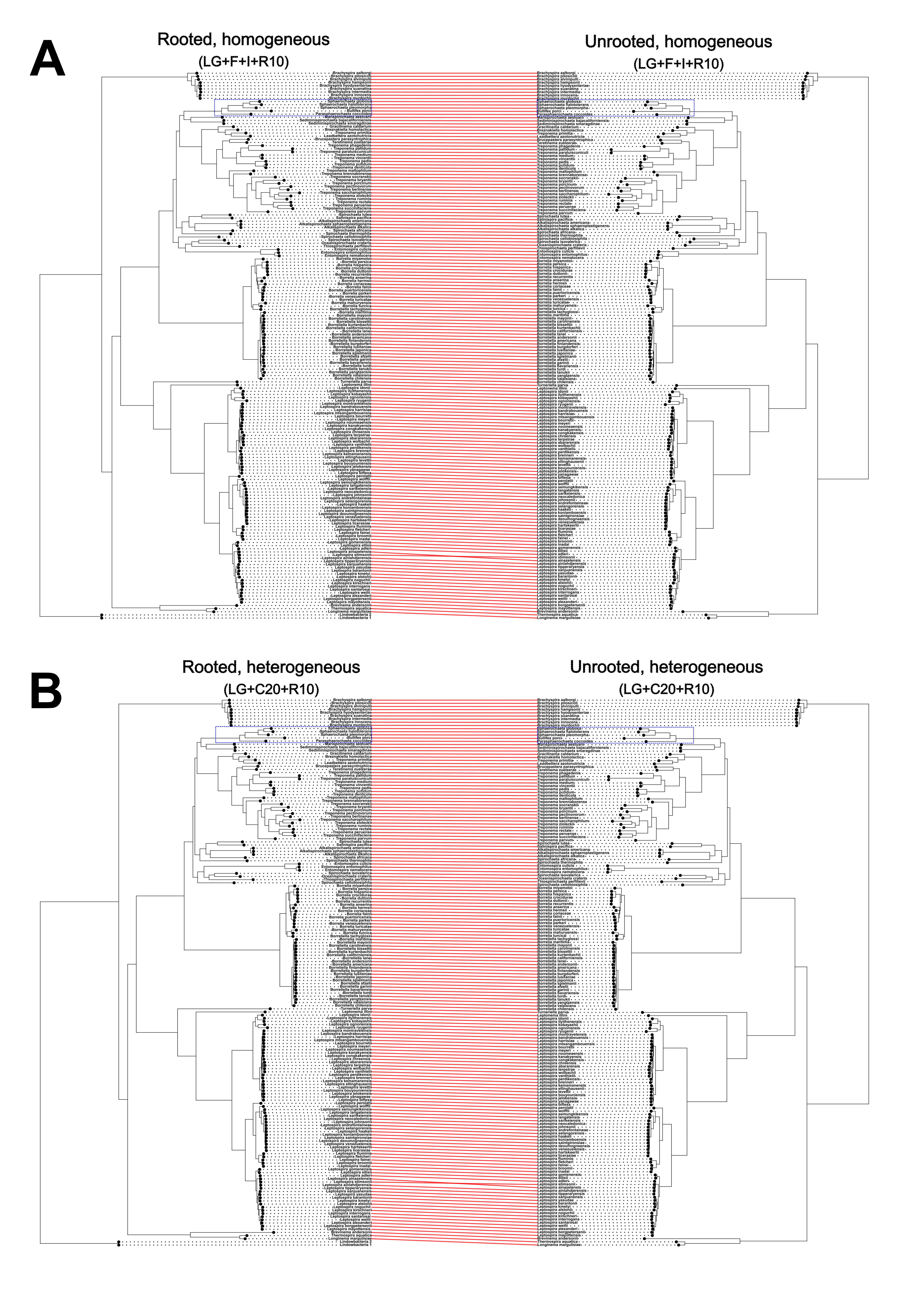

### FigS4

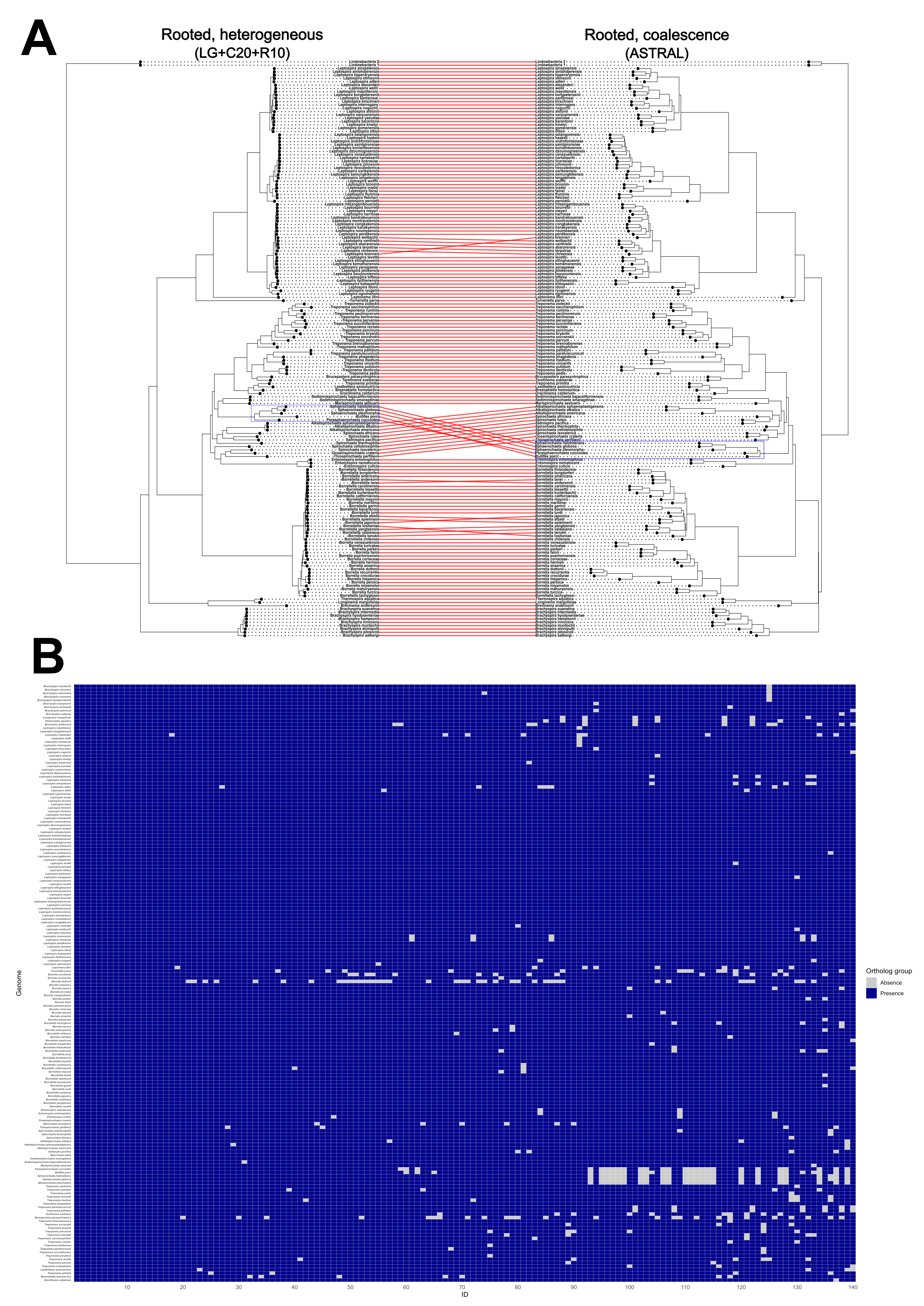

### FigS5

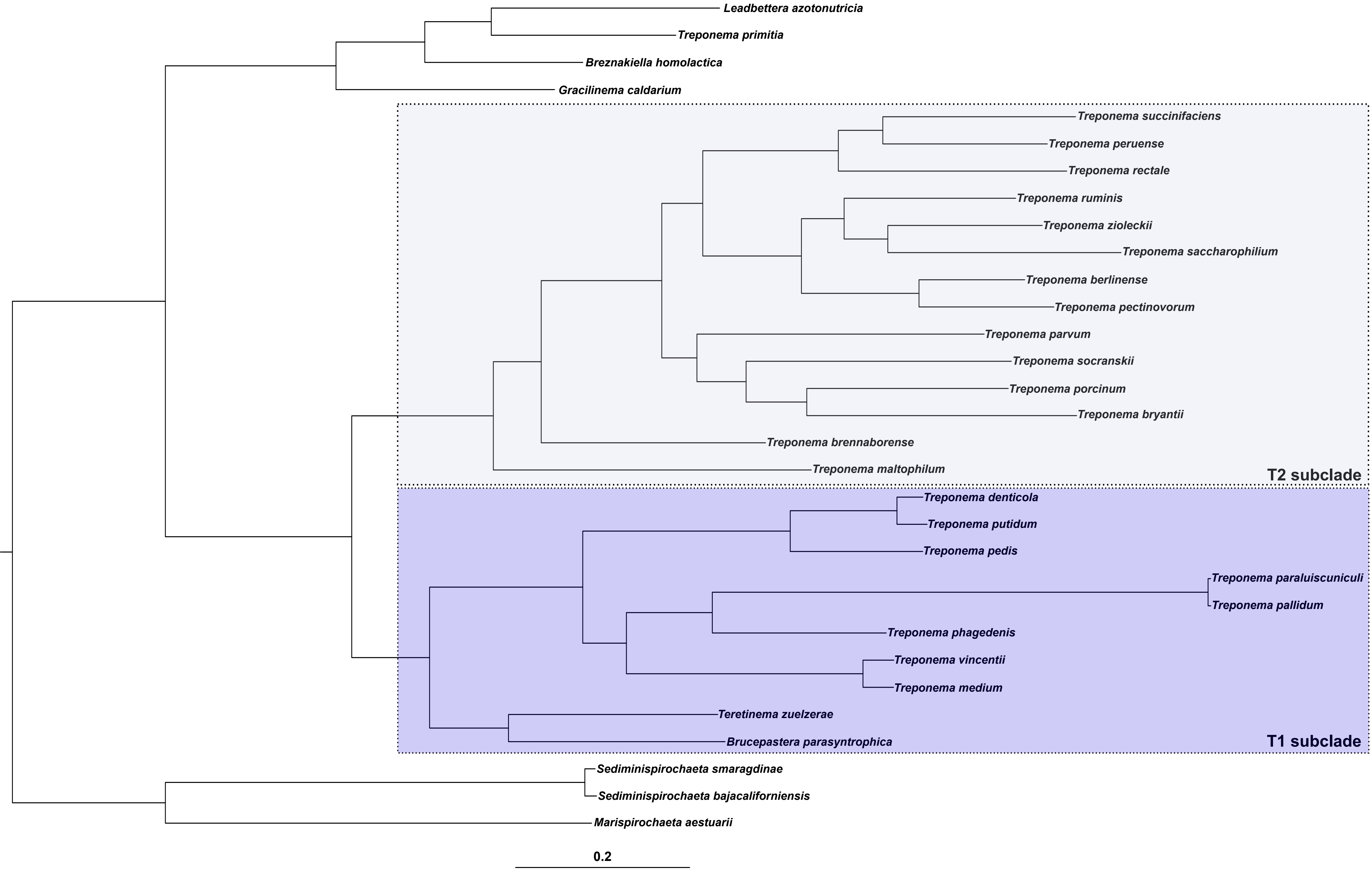

### FigS6

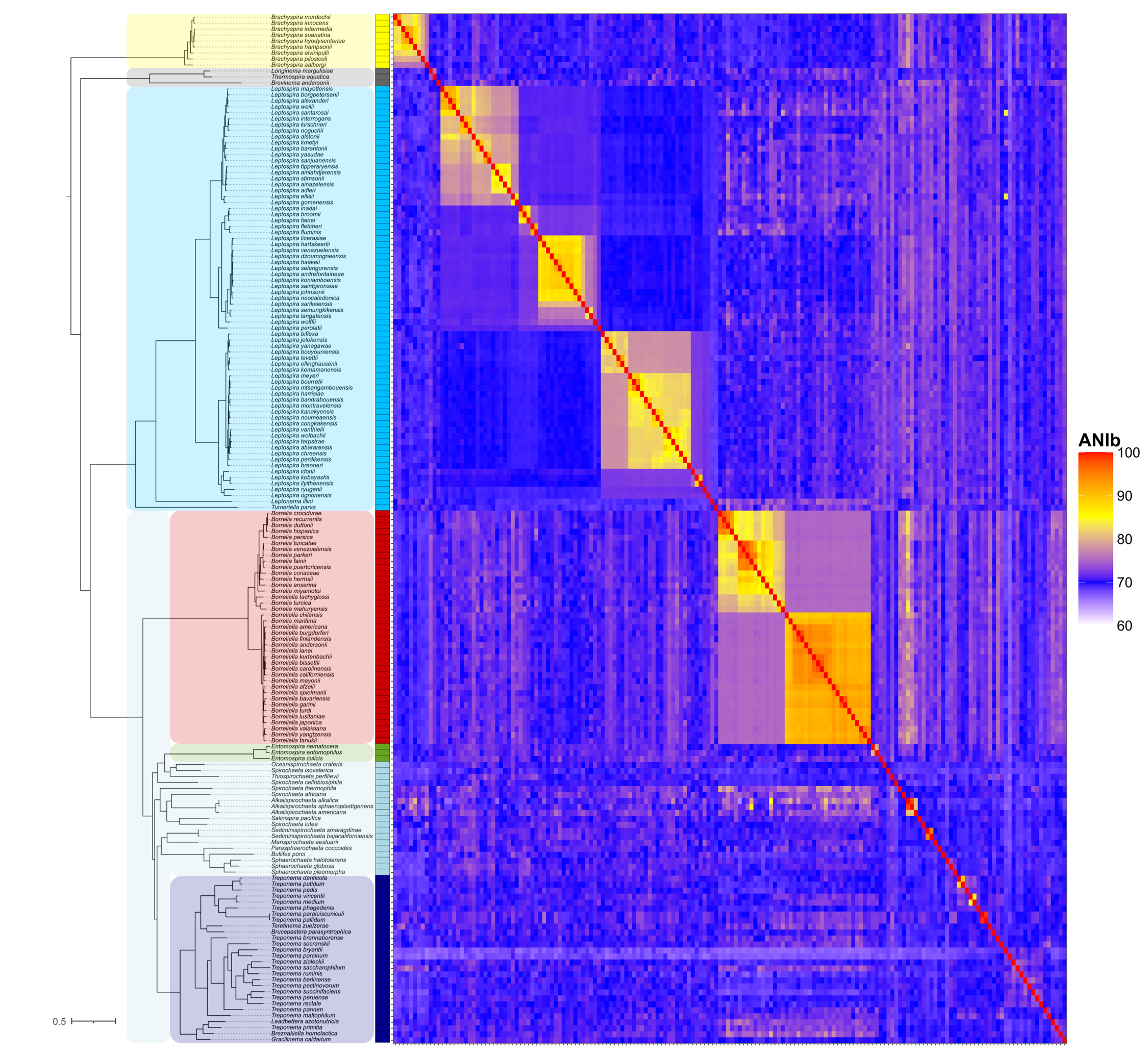

### FigS7

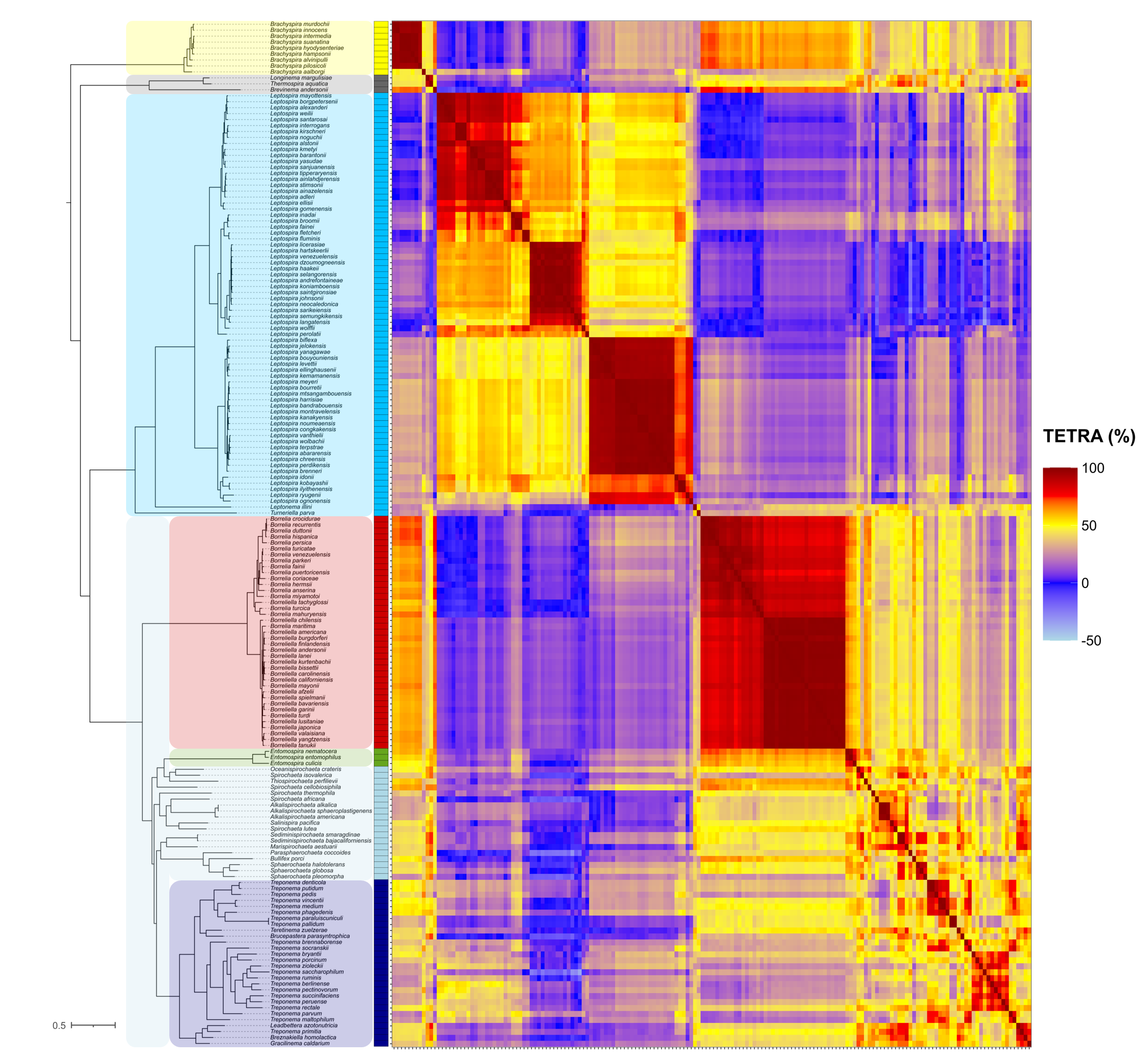

### FigS8

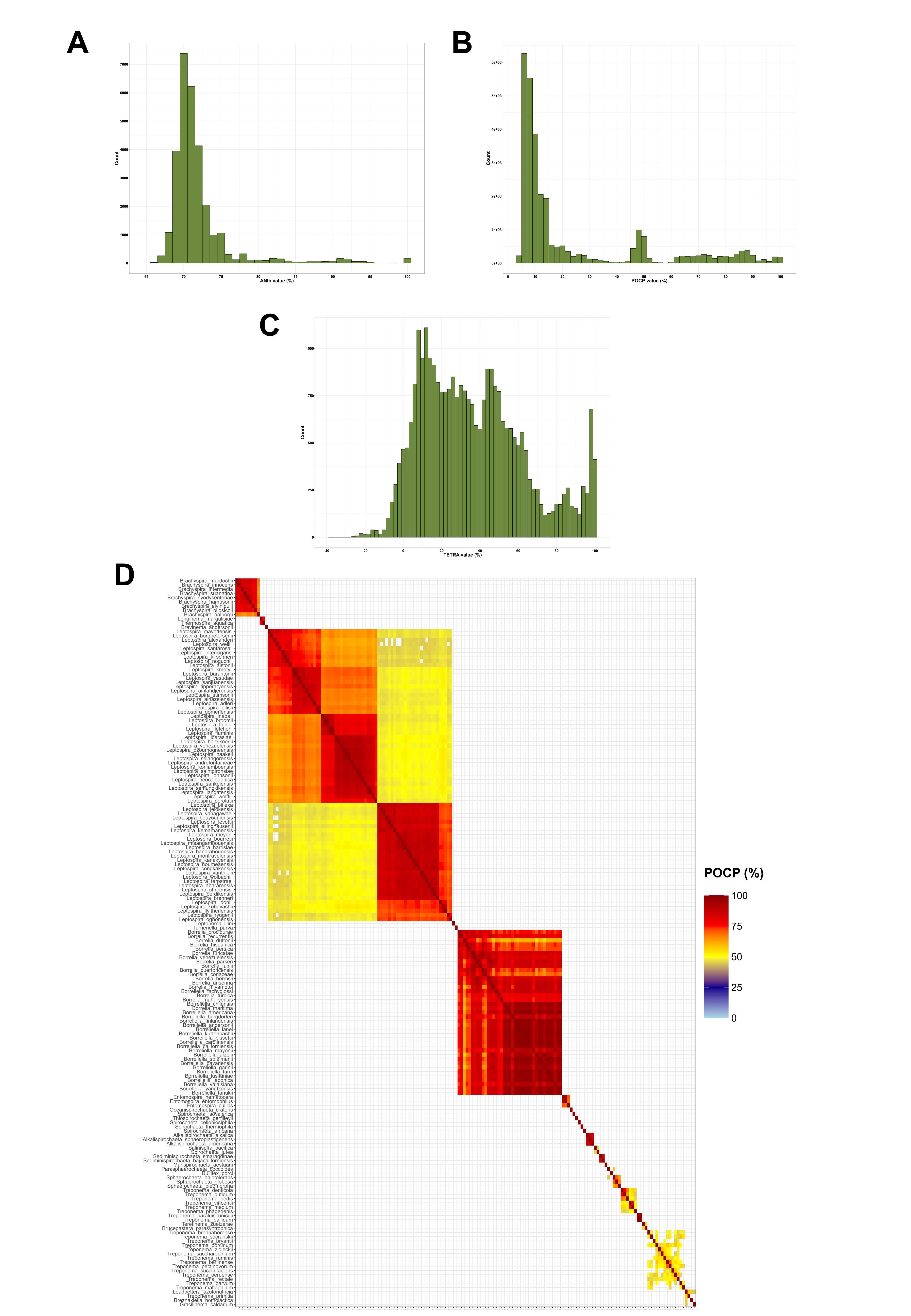

### FigS9

**A**

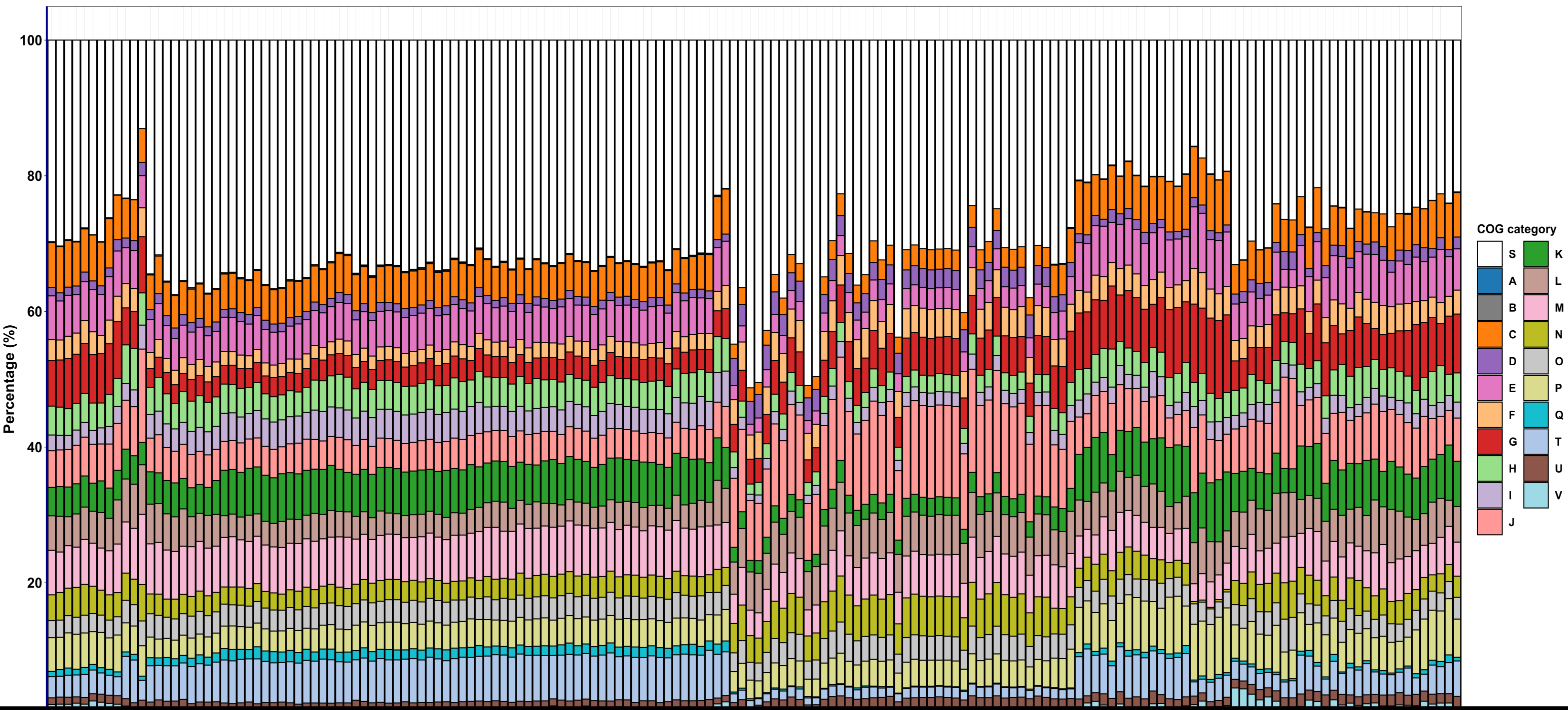


[redacted]

**B**

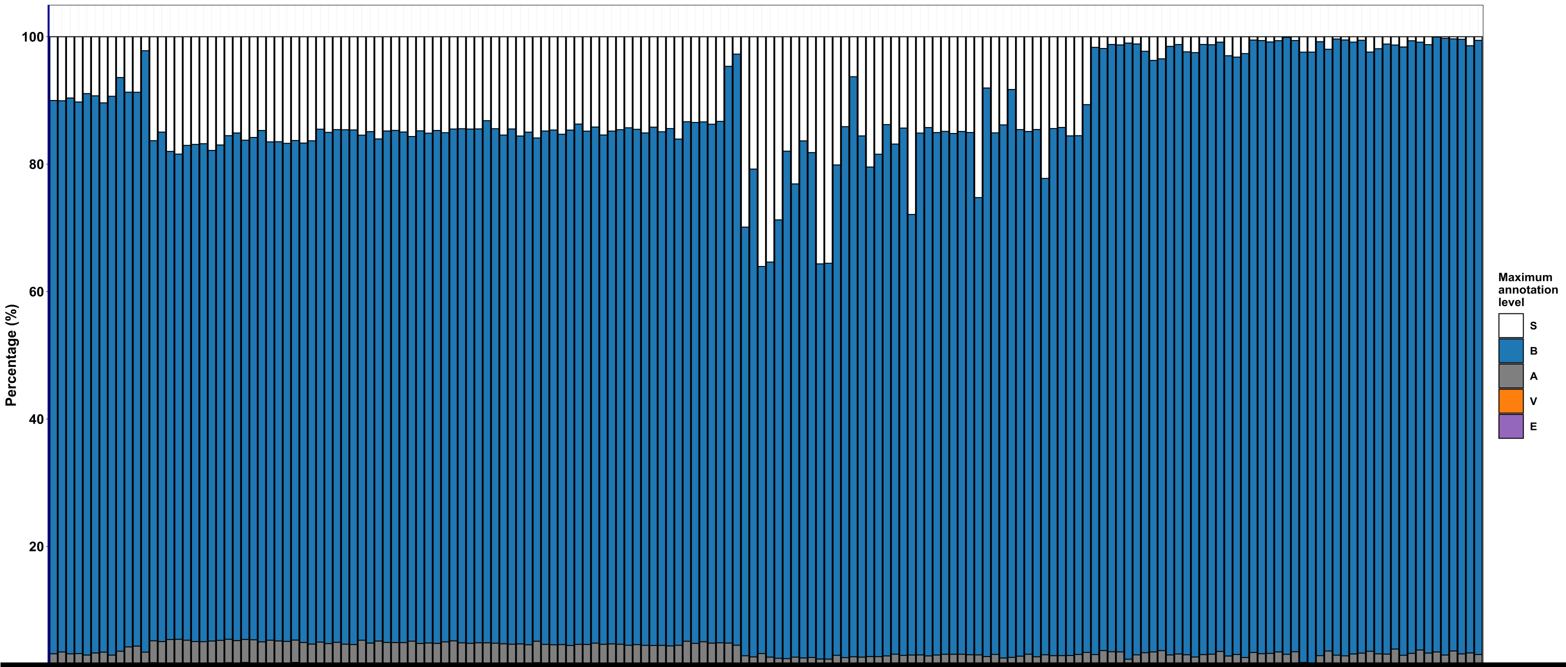


[redacted]

### FigS10

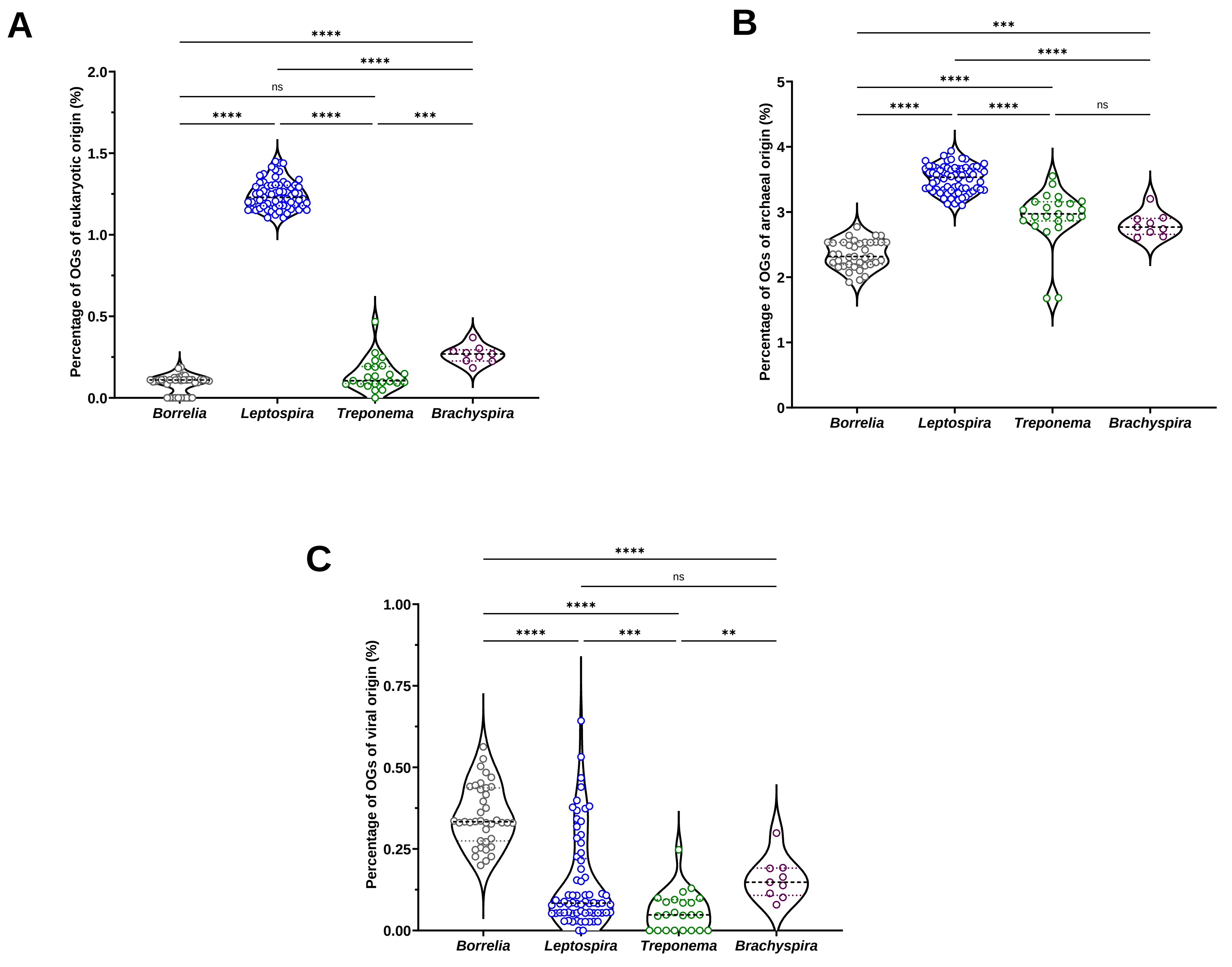

### FigS11

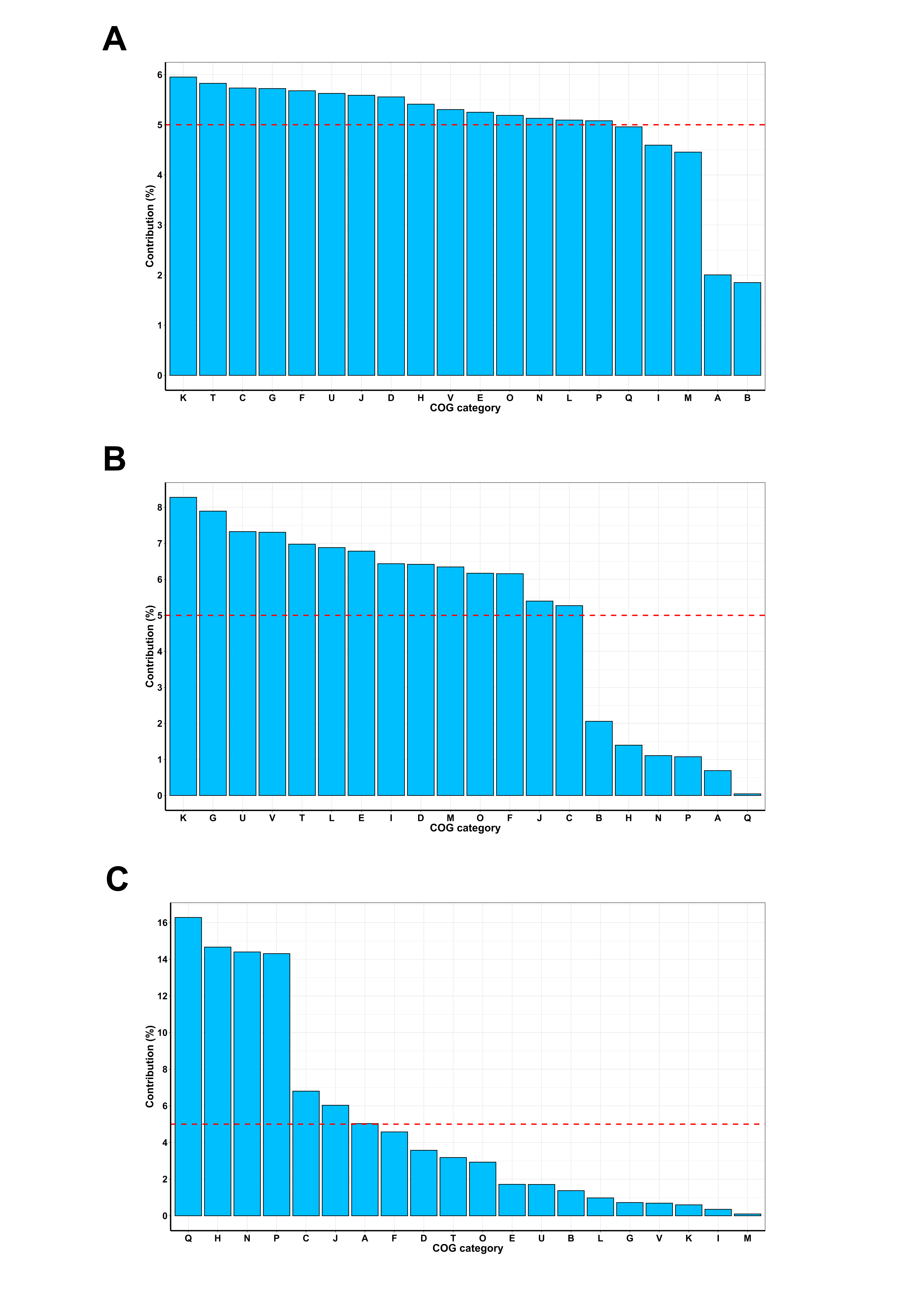

### FigS12

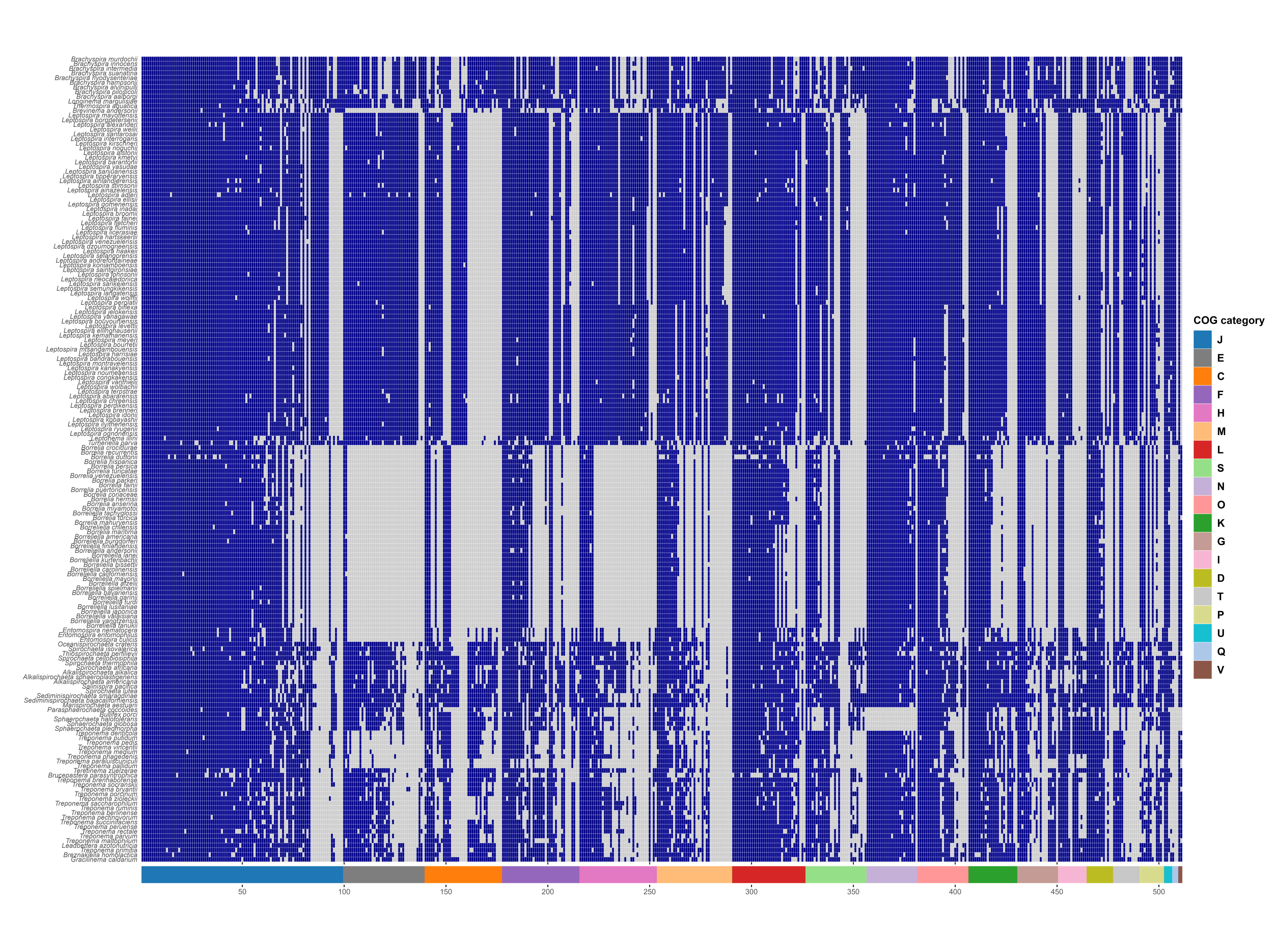

### FigS13

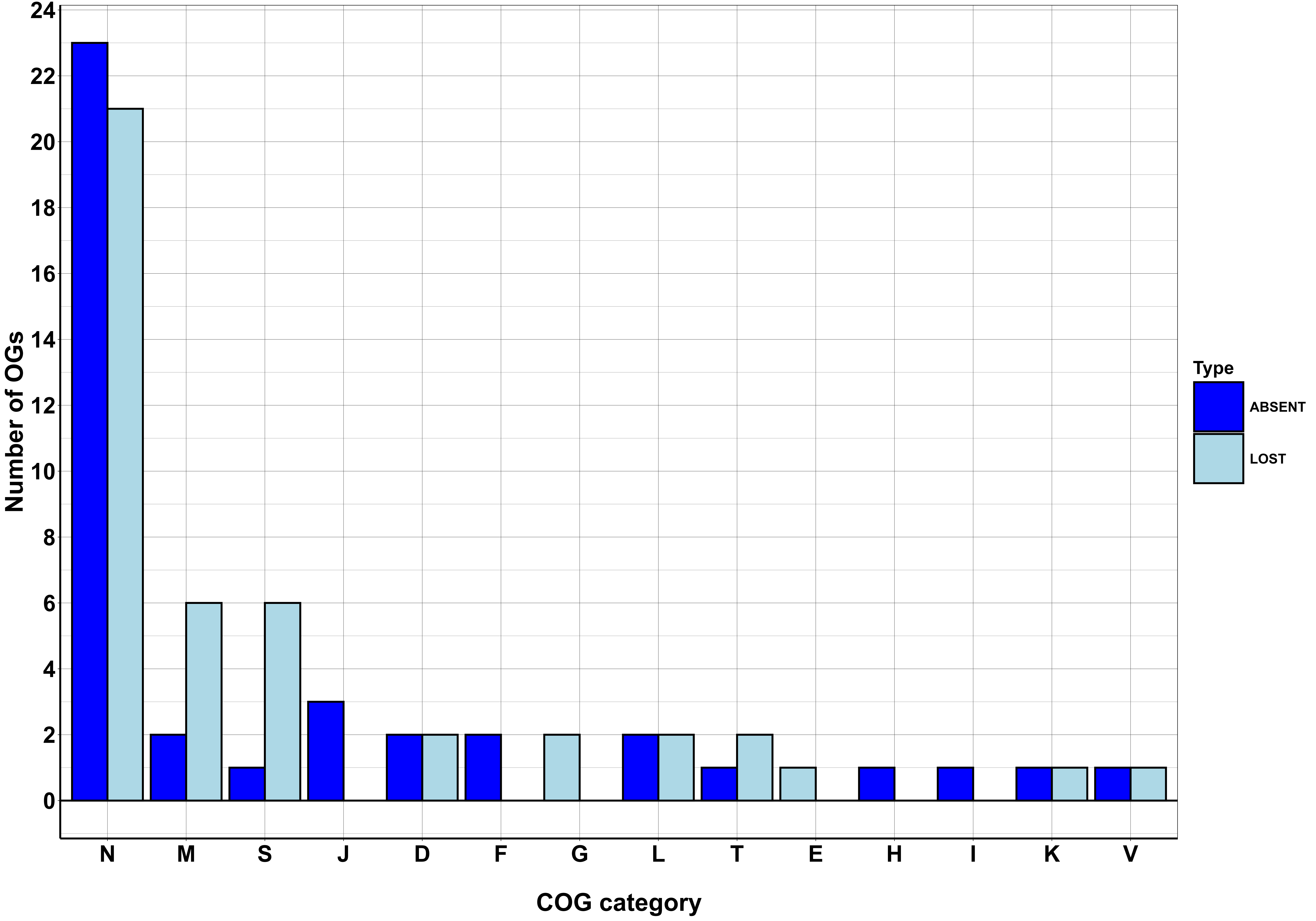
